# Oncogene-induced senescence and senescence-associated secretory phenotype (SASP) determine the efficacy of palbociclib in PIK3CA mutated colorectal cancer

**DOI:** 10.1101/2024.11.15.623304

**Authors:** Pingping Cao, Xiaohui Zhou, Suisui Yang, Tianqi Shen, Shuai Wang, Hanyang Yu, Xiaorong Liu, Yeqing Gong, WenHong Wang, Haiyang Wang, Tingting Zhou, Jing Wang, Zhining Fan, Mingde Huang, Xu Qian, Xiuxing Wang, Qianghu Wang, Liu Yang, Yingjian Zhang, Fan Lin

## Abstract

Palbociclib is an excellent CDK4/6 inhibitor, but its clinical application is mainly limited to the treatment of advanced ER^+^/HR^+^ and HER2^−^ breast cancer. Its efficacy in colorectal cancer (CRC) is evaluated in multiple trials and so far remains undetermined. We found that the PIK3CA-mutant CRC cells were insensitive to palbociclib treatment compared with the PIK3CA wild-type counterparts, and they were in an OIS (oncogene-induced senescence) state which was predisposed to a strong senescence-associated secretory phenotype (SASP) upon treatment. These senescent cells excessively secreted various SASP factors LCN2, and ultimately caused palbociclib-resistance via upregulation of EGFR in the non-senescent CRC cells. Importantly, a drug combination screen identified that erlotinib could synergize with palbociclib to overcome the SASP-induced resistance. Overall, we found that PIK3CA mutation-induced senescence compromised the efficacy of palbociclib treatment in CRC but co-targeting EGFR could minimize the OIS-associated side effects while preserving the beneficial effects.

## INTRODUCTION

Colorectal cancer (CRC) is a global leading cause of cancer-related morbidity and mortality in both men and women ^1^. Currently, radio– and chemo-therapy are still the mainstays for post-surgical treatment of CRC, and are often accompanied by severe side effects, limited efficacy, frequent recurrence, and metastasis ^2^. Palbociclib (Ibrance™, Pfizer) is the first CDK4/6 inhibitor that has been approved by the FDA in 2015 for the first-line treatment of postmenopausal women with HR/ER-positive and HER2-negative advanced breast cancer in combination with letrozole ^3^. Given its good clinical efficacy in breast cancer, CDK4/6 inhibitors are being tested in several clinical trials for CRC, both alone and with chemotherapeutic/targeted agents (https://clinicaltrials.gov). Despite some progress being made, the effectiveness of palbociclib for CRC treatment remains uncertain, and further investigations are still required ^4, 5^.

Abnormal cell proliferation caused by dysregulation of the cell cycle is one of the hallmark characteristics of tumors ^6^. Normally, gradual accumulation of cyclin in the G1 phase activates CDK4 and/or CDK6, two Cyclin-dependent kinases, leading to phosphorylation of the RB and activation of E2F transcription of downstream genes required by cell cycle progression ^7^. Whereas, CDK4/6 are often overexpressed or continuously being activated in various tumors including CRC cells, so their cell cycles are often accelerated ^8^. Palbociclib or several other CDK4/6 targeted inhibitors exert their activities by competitively binding to the ATP-binding pocket of CDK4/6 to inactivate the CDK4/6-Cyclin D complexes, leading to inhibition of RB phosphorylation and subsequent blockage of the cell cycle.

Besides the cell cycle arrest, another therapeutic effect caused by palbociclib is the induction of senescence ^9–11^. Cellular senescence is a state of terminal growth arrest in which cells are unresponsive to growth factor stimulation, and it could be induced by telomere shortening, intracellular ROS accumulation, oncogene activation, chemo-or radiotherapy, and cell cycle targeted therapy in our case ^12–15^. Senescent cells permanently lose their ability to re-enter the cell cycle and proliferate, so both oncogene-induced senescence (OIS) and therapy-induced senescence (TIS) are regarded as a protective mechanism or treatment-favorable roles for preventing tumor genesis and development ^16^.

However, in this study, we found a novel phenomenon that the OIS caused by the gain-of-function mutation of PIK3CA may not act as a protective barrier but as an accomplice of palbociclib resistance and tumor progression. In these cells, we not only observed strong expression of senescence markers which was putatively driven by OIS, but also found that the senescence phenotype was further aggravated by palbociclib treatment. Unfortunately, the additive effect of palbociclib-induced TIS and PIK3CA-mutation-induced OIS was found to prominently counteract the efficacy of palbociclib, but not the opposite.

Senescent cells have pleiotropic functions in cancer, aging, and other diseases by secreting inflammatory cytokines, chemokines, and matrix proteases (known as the senescence-associated secretory phenotype, SASP), and cause beneficial or detrimental effects on the tumor or its microenvironment in a context-dependent manner ^17, 18^. In the present study, we discovered that the SASP factor(s) secreted by the PIK3CA mutant CRC cells conferred resistance to palbociclib via upregulation of EGFR in the adjacent non-senescent cells, and it could be overcome by combining palbociclib with an EGFR inhibitor. In this way, we proposed a novel senotherapy that only targets the major effector molecule instead of trying to eliminate all the senescent cells in the senolytic therapy or suppress the SASP in the senomorphic therapy, and the beneficial side of palbociclib-induced senescence could be preserved.

## RESULTS

### 1. Gain-of-function mutations of the PIK3CA gene confer resistance to palbociclib in colorectal cancer cells

Firstly, we assessed the anticancer efficacy of palbociclib in CRC cell lines and accidentally found a tendency that those carrying gain-of-function mutation of the PIK3CA gene were more resistant to palbociclib treatment in both anti-proliferation (Fig. 1A) and clonogenic experiments (Fig. 1B). The average IC50 of three PIK3CA mutant CRC cell lines, namely, HCT116, HCT15, and DLD1 were approximately 2-fold higher than that of LOVO and SW620, two CRC cell lines carrying wild-type PIK3CA (Fig. 1A and C). In addition, the clinically relevant plasma concentration of palbociclib (up to 500 ng/mL ≈ 1.12 µM) ^19^ failed to cause sufficient apoptosis in PIK3CA mutant cell lines (Fig. S1A and B). Interestingly, this is not due to the insufficient inhibition of the targeted protein or arrest of the cell cycle by palbociclib treatment, as the level of p-RB and FoxM1 were markedly declined from 0.5 µM related to control (Fig. 1D), and substantial cells were detained in G1 phase even at a very low concentration of 0.1 µM in PIK3CA mutant cell lines (Fig. S1D and E). By a joint analysis of the drug sensitivity data from GDSC (https://www.cancerrxgene.org/) and mutation data of CRC cell lines from COSMIC (http://cancer.sanger.ac.uk), only TP53 and PIK3CA exhibited much higher mutation frequency in palbociclib resistant (27 cell lines) than sensitive (14 cell lines) CRC cell lines in the top 10 most frequently mutated genes in all CRC cell lines (Fig. 1E). Nevertheless, the association between the palbociclib sensitivity and the genetic status of TP53 was not as strong as PIK3CA in our CRC cell lines (Fig. S1C). Moreover, PIK3CA is the 5th most frequently mutated gene in 386 clinical CRC samples and the missense mutations of E545K/G and H1047R/L are two predominant forms of its mutation which also occurred in HCT116 (H1047R/G), HCT15 and DLD1 (E545K/L) cell lines (Fig. 1F).

**Figure 1.**
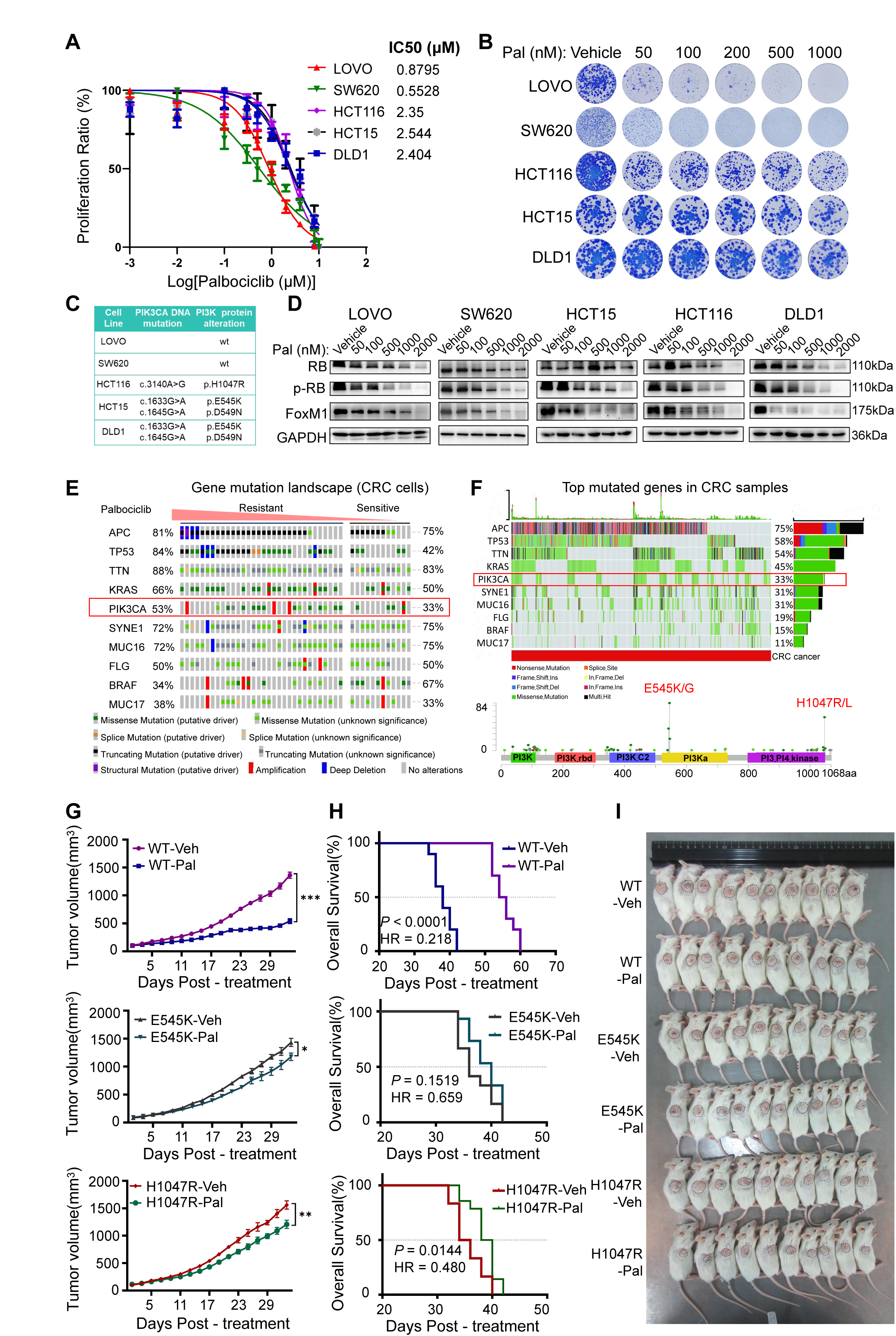
Palbociclib antitumor activity and PIK3CA mutations promote palbociclib resistance in colorectal cancer. **A**, Growth inhibition curves and IC50 values for five CRC cell lines treated with palbociclib. **B,** Images showing the colony formation of five CRC cell lines after palbociclib treatment. **C,** Table of PIK3CA gene mutations in five CRC cell lines from the CCLE database. **D,** Immunoblot for RB, p-RB and FoxM1 from five CRC cell lines with palbociclib treatment. **E,** Analysis of the mutational profiles of palbociclib-resistant (IC50 > 10 μM) and sensitive (IC50 < 10 μM) cells from the GDASC database. **F,** Mutational landscape of CRC from TCGA data and PIK3CA gene mutations (E545K/G and H1047R/L) from the GSCA database. **G,** Tumor volume curves of CDX models (LOVO-PIK3CA WT and MUT cells) after vehicle or palbociclib treatment. **H,** Kaplan-Meier curves depicting OS for CDX mice (LOVO-PIK3CA WT and MUT cells) treated with vehicle or palbociclib. **I,** Representative images of CDX mice (LOVO-PIK3CA WT and MUT cells) after vehicle or palbociclib treatment. Mean ± SD; Mean ± SEM; *, *P* < 0.05, **, *P* < 0.01; and ***, *P* < 0.001.

If the therapeutic effect of palbociclib in the treatment of CRC was indeed determined by the genetic status of the PIK3CA gene, the clinical efficacy of CDK4/6 targeted therapy could be compromised in about 1/3 of all CRC patients. To confirm this finding, we constructed PIK3CA wild-type (WT) and mutant (MUT) isogeneic cell lines using LOVO cells that originally carried the WT PIK3CA gene. The levels of phosphorylated AKT of PIK3CA E545K (2# and 5# clones) and H1047R (2# and 3# clones) cells were much higher than the WT counterparts, confirming their PI3K-AKT signaling being continuously activated (Fig. S1F). Moreover, E545K or H1047R cells were more resistant to palbociclib than the WT control in the anti-proliferation (Fig. S1G) and colony formation experiments (Fig. S1H).

Next, we established a xenograft model by subcutaneously injecting the aforementioned WT, E545K (2#), and H1047R (3#) LOVO cells to immunodeficient NNG mice to further validate the PIK3CA-associated resistant phenotype *in vivo*. As shown in Fig. 1G-I, oral administration of 25 mg/kg palbociclib caused significant tumor suppression and prolonged overall survival in mice carrying WT PIK3CA xenograft. In contrast, palbociclib only marginally suppressed the growth of PIK3CA MUT tumors and failed to cause a statistically meaningful improvement in the survival of these mice. The H&E and Ki-67 staining of tumor tissue also reflected a similar trend (Fig. S1I). Lastly, we did not observe marked body weight loss or any other severe side effects of mice in all groups during the three-week treatment (Fig. S1J).

### 2. Palbociclib treatment aggravated the PIK3CA mutation-induced senescence and senescence-associated secretory phenotype

To interrogate the molecular mechanism that PIK3CA mutation conferred resistance to palbociclib, we performed RNA-seq for PIK3CA WT/MUT LOVO cells treated with palbociclib or vehicle solution (Fig. 2A). KEGG pathway analysis revealed that Cellular Senescence was the top 6th enriched pathways in WT cells (Fig. 2B), while it was ranked the top 3rd among all enriched pathways in MUT cells upon palbociclib treatment, suggesting that the post-treatment MUT cells possessing a stronger senescence profile than WT cells (Fig. 2C). β-galactosidase staining and Western blotting results confirmed that palbociclib treatment rendered a stronger senescence phenotype in PIK3CA MUT cells than WT cells (Fig. 2D-F, Fig. S2A and B). Notably, untreated MUT and WT cells had similar levels of cell cycle-related proteins, but MUT cells exhibited higher baseline levels of p21, p27, and p53 than WT cells (Fig. 2G-H).

**Figure 2.**
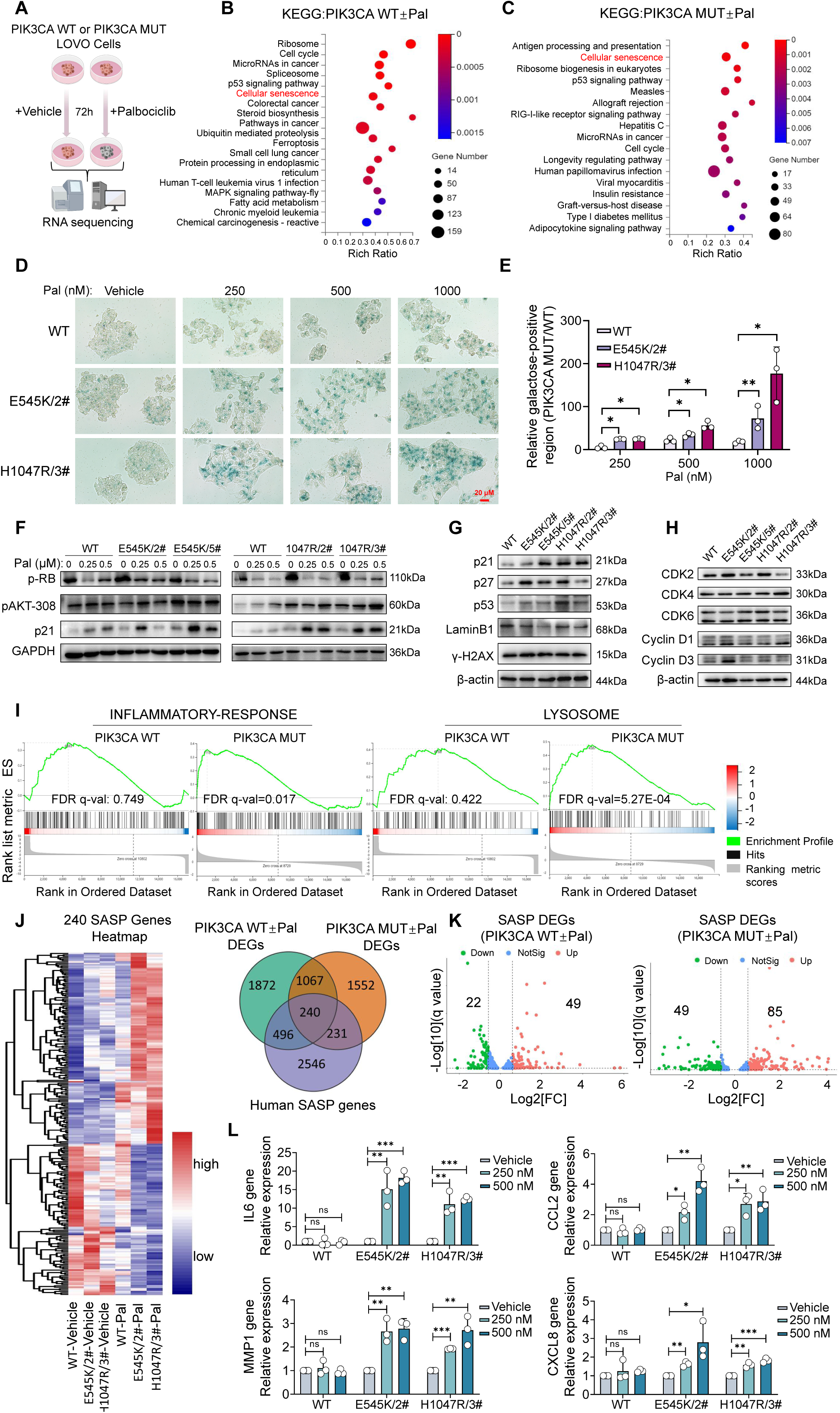
PIK3CA mutations enhance palbociclib-induced senescence and alter SASP gene expression. **A,** Schematic of RNA-sequencing on LOVO-PIK3CA WT and MUT cell lines treated with palbociclib. KEGG analysis of RNA-sequencing datas from LOVO-PIK3CA WT (**B**) and MUT (**C**) cell lines with palbociclib treatment. **D,** β-galactosidase staining of LOVO-PIK3CA WT and MUT cell lines treated with palbociclib (Scale bar: 20 μm). **E,** Quantitation of the percentage of β-galactosidase-positive cells (by image **D**). **F,** Immunoblot for p-RB, p21, pAKT-308 from LOVO-PIK3CA WT and MUT cell lines with palbociclib treatment. **G,** Immunoblot analysis of Senescence marker proteins in LOVO-PIK3CA WT and MUT cell lines. **H,** Immunoblot of G1 phase regulatory proteins from LOVO-PIK3CA WT and MUT cell lines. **I,** GSEA enrichment analysis of RNA-sequencing data from LOVO-PIK3CA WT and MUT cell lines with palbociclib treatment. **J,** Venn diagram (right) depicting overlap between SASP genes in palbociclib-treated LOVO-PIK3CA WT and MUT cell lines. Heatmap (left) depicting the expression of 240 common SASP genes. **K,** Volcano plot of SASP genes expression from RNA-sequencing datas. **I,** mRNA expression of SASP genes from LOVO-PIK3CA WT and MUT cell lines treated with palbociclib. Mean ± SD; *, *P* < 0.05, **, *P* < 0.01; and ***, *P* < 0.001.

Meanwhile, GSEA revealed that palbociclib treatment resulted in significant enrichment of the Inflammatory Responses and Lysosomes pathways genes (*p* = 0.017 and *p* = 5.27E-04, respectively) in MUT but not in WT cells (Fig. 2I). Since both lysosomal dysfunction and aberrant local inflammation were features of cellular senescence ^20–22^, we postulated that the upregulation of inflammatory response pathways in palbociclib-treated MUT cells may be related to the alteration of cytokine/chemokine-factors secreted caused by the SASP ^23^. As shown in Fig. 2J Venn diagram, palbociclib treatment resulted in 496 and 231 exclusive differentiated expressed genes (DEGs) of SASP factors (SASPF) in WT and MUT cells, respectively, and 240 common SASPF-DEGs in both WT and MUT cells. Heatmap analysis of the 240 common SASPF-DEGs also showed that the expression patterns of the vehicle-treated WT and MUT cells were similar, whereas those of palbociclib-treated MUT cells were much different from the PIK3CA WT cells (Fig. 2J Heatmap). Moreover, 85 SASPF genes were significantly upregulated (log2FC >1.5) in MUT cells, compared to only 49 in WT cells treated with palbociclib (Fig. 2K). To confirm the pronounced alteration of SASPF expression in MUT cells, q-PCR was used to assess the changes of several representative SASPF genes with/without treatment. As shown in Fig. 2L, palbociclib treatment significantly elevated the expression of IL6, CCL2, MMP1, and CXCL8 in MUT but not in WT LOVO cells, and this phenomenon was also observed in the CRC cell lines carrying intrinsic PIK3CA mutation (Fig. S2C and D).

### 3. SASP factors secreted from palbociclib-treated PIK3CA mutant cells caused resistance to palbociclib via elevation of EGFR

We wondered whether the altered expression of SASPF genes in palbociclib-treated MUT cells could be the cause of palbociclib resistance, so a series of experiments were designed to evaluate the effect of the condition medium collected from vehicle-treated WT/MUT LOVO cells (Control CM) or palbociclib-induced senescent WT/MUT cells (S-CM) (Fig. 3A). Compared with the post-treatment S-CM derived from WT cells, the MUT cells derived S-CM was capable of accelerating the short-term proliferation (Fig. 3B), long-term colony formation (Fig. 3C), and conferring resistance to palbociclib in the same type of cells cultured with S-CM (Fig. 3D).

**Figure 3.**
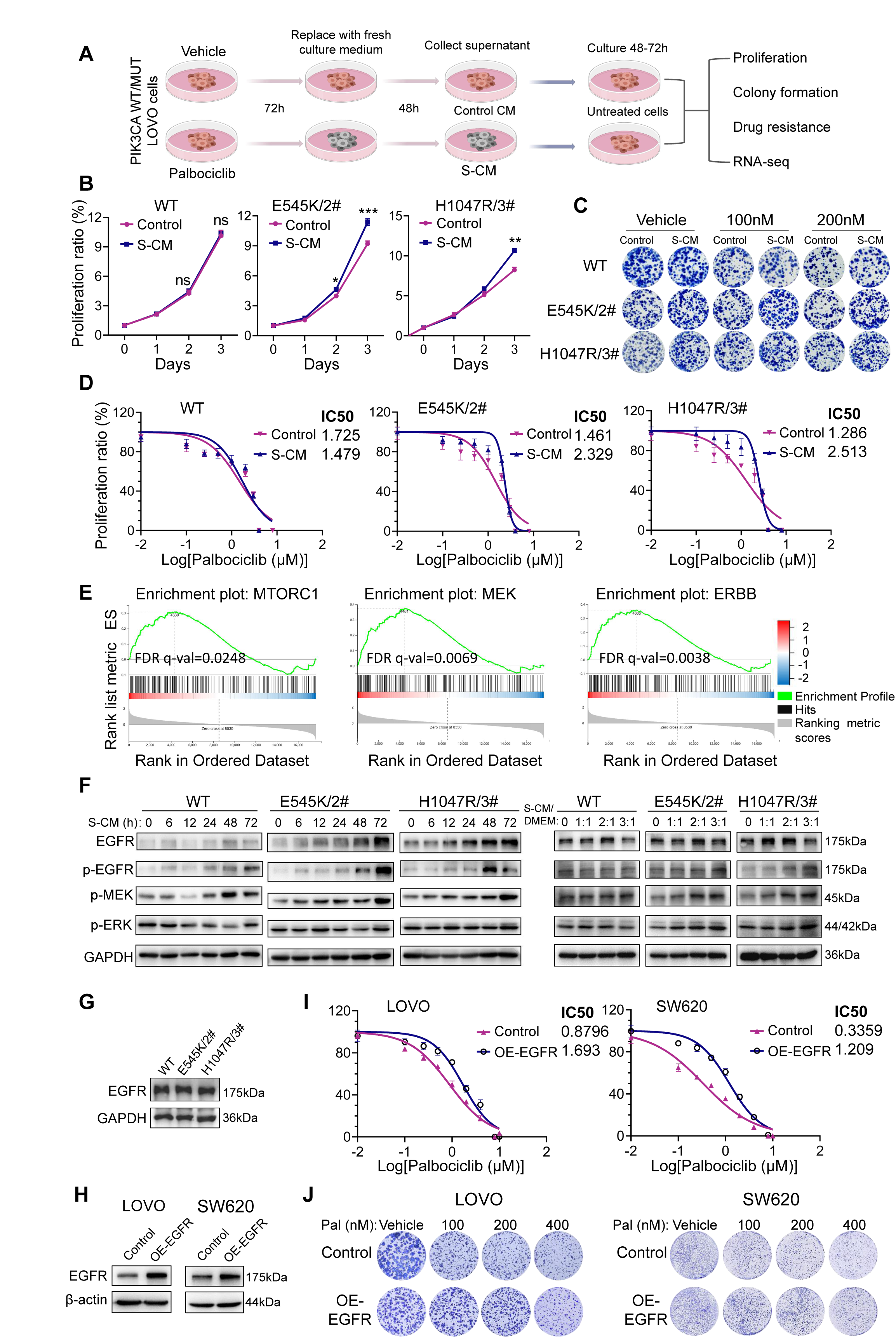
SASP factors in palbociclib-induced PIK3CA mutant cell lines lead to resistance by upregulating EGFR. **A,** Schematic illustrating the experimental approach of treating LOVO-PIK3CA WT and MUT cell lines with palbociclib, then collecting senescent conditioned medium (S-CM) for further experiments. **B,** Short-term proliferation assays of LOVO-PIK3CA WT and MUT cell lines upon treatment S-CM and control. **C,** Images showing the colony formation of LOVO-PIK3CA WT and MUT cell lines upon treatment S-CM and control. **D,** Growth inhibition curves and IC50 values of LOVO-PIK3CA WT and MUT cell lines exposed to S-CM and control under palbociclib treatment. **E,** GSEA enrichment analysis of RNA-sequencing datas from LOVO-PIK3CA MUT cell lines with S-CM treatment. **F,** Immunoblot for EGFR pathway protein expression in LOVO-PIK3CA WT and MUT cell lines under S-CM time (0, 12, 24, 48, and 72 hours) and concentration (S-CM S-CM/DMEM = 1:1, 2:1, and 3:1) gradient treatment. **G,** Immunoblot for EGFR expression in LOVO-PIK3CA WT and MUT cell lines. **I,** Growth inhibition curves and IC50 values for LOVO and SW620 cells overexpressing EGFR under palbociclib treatment. **H,** Immunoblot for EGFR expression in LOVO and SW620 cells overexpressing EGFR gene. **J,** Images showing the colony formation of LOVO and SW620 cells after overexpression of EGFR genes with palbociclib treatment. Mean ± SD; *, *P* < 0.05, **, *P* < 0.01; and ***, *P* < 0.001.

To determine which pathways might be involved in the S-CM-induced palbociclib resistance, KEGG and GSEA analyses of the RNA-seq data from Control CM *vs* S-CM-treated MUT cells were performed. As shown in Fig. 3E and Fig. S3A, S-CM treatment caused significant enrichment of the PI3K-AKT/mTOR and MAPK/MEK signaling pathways. Moreover, the ERBB family-related gene sets were also significantly enriched (Fig. 3E). Therefore, we inferred that the aberrant EGFR and its downstream signaling could be the cause of palbociclib resistance. To confirm this finding, the WT/MUT LOVO cells were treated with S-CM for different time durations or dilution ratios. As a result, S-CM caused elevations of total EGFR and p-EGFR, and its downstream molecules in both WT and MUT cells over time, while the elevations were more prominent in the MUT than WT cells (Fig. 3F). Meanwhile, increasing the mixing ratio of S-CM/DMEM also caused elevation of EGFR and p-EGFR in MUT cells. Importantly, the basal level of EGFR between the untreated WT and MUT cells was not different (Fig. 3G), so the upregulation of EGFR was not directly caused by PIK3CA mutation. Moreover, immunohistochemical staining revealed that palbociclib treatment (25 mg/kg, p.o.) significantly increased EGFR expression in MUT tumors from xenograft mice, while such increase was not seen in WT tumors (Fig. S3F and G).

The next question was whether the elevated EGFR level responsible for the palbociclib resistance. To study that, we established EGFR overexpressed (OE-EGFR) LOVO and SW620 cells (Fig. 3H) for quantitative proteome analysis (DIA-MS). Overexpression of EGFR markedly increased the tolerance of LOVO and SW620 cells to palbociclib treatment in the anti-proliferation (Fig. 3I) and colony formation (Fig. 3J) experiments. GO analysis of the proteomic data discovered that the EGFR overexpression resulted in significant enrichment in cell division, cell cycle, G2/M transition, and mitosis-related pathways (Fig. S3C). In addition, GSEA revealed cell cycle-related genes were enriched in both S-CM treated MUT (Fig. S3B) and OE-EGFR LOVO cells (Fig. S3D), suggesting similar effects from S-CM treatment or EGFR overexpression. The Heatmap showed a distinct expression pattern of cell cycle-related genes in OE-EGFR cells compared to parental LOVO cells (Fig. S3E). Notably, some key cell cycle genes like CDK2, RB1, MDM2, and WEE1, which are significantly altered in OE-EGFR cells, have been reported to cause palbociclib resistance. To obtain a comprehensive profile of EGFR-induced palbociclib resistance, we compiled a palbociclib-resistance gene set by merging patient-derived and literature-based palbociclib resistance genes. (Fig. S3H and Supplemental Table 1) ^24^. As a result, the GSEA discovered that these palbociclib resistance genes were significantly enriched in both S-CM-treated MUT and OE-EGFR LOVO cells (Fig. S3I). Finally, WB confirmed several representative cell cycle-related proteins and proteins that were involved in palbociclib resistance (Fig. S3J) in OE-EGFR LOVO cells (Fig. S3K). These results indicated that S-CM induced EGFR overexpression, leading to extensive changes of multiple genes that collectively enhanced the palbociclib resistance in recipient CRC cells.

### 4. SASP factor LCN2-mediated EGFR upregulation was essential for palbociclib resistance

To identify which factor upregulating EGFR in S-CM, we examined SASPF genes markedly upregulated (> 2-fold) in palbociclib-treated PIK3CA MUT cells. Of the 32 SASPF genes commonly upregulated in both WT and MUT cells, 16 showed higher expression in MUT cells (Fig. S4A). Additionally, 53 out of 85 significantly upregulated SASPF genes were exclusively found in palbociclib-treated MUT cells but not in WT cells. The differentiated gene expression pattern of the total 69 selected SASPF genes in vehicle/palbociclib-treated MUT cells was illustrated in a heatmap in Fig. 4A. Next, we analyzed the potential interactions between 69 SASPF and EGFR using the STRING database, finding 23 SASPF have interactions with EGFR (Fig. S4B), with 12 showing strong interactions (combined-score >0.7, Fig. S4C). One of them is LCN2, a gene encoding lipocalin-2 and being reported to interact with EGFR to facilitate its recycling under TGF-α stimulation ^25^. Q-PCR and ELISA assays revealed that palbociclib treatment significantly increased LCN2 expression and secretion in MUT cells while not affecting WT cells. (Fig. 4B and C). Importantly, overexpression of LCN2 in LOVO and SW620 cells caused elevation of EGFR in the protein level (Fig. 4D and E), but not in the mRNA level (Fig. 4F and G). Moreover, correlation analyses of CRC cell lines from CCLE showed a strong positive correlation between EGFR and LCN2 protein levels (R = 0.57, *P* = 0.02) and their correlation at the mRNA level was less prominent (R = 0.29, *P* = 0.03, Fig. S4D). Lastly, LCN2 overexpressed LOVO and SW620 cells were more resistant to palbociclib than their parental cells in colony formation (Fig. 4H and I). Based on the above evidence, we reckoned that LCN2 may regulate EGFR via the protein-protein interaction. Indeed, confocal microscopy showed significant colocalization of EGFR (red) and LCN2 (green) in CRC cells overexpressing exogenous LCN2 and EGFR, and the two immunofluorescent signals were highly overlapped (Fig. 4J and K). Furthermore, exogenous EGFR-Flag/EGFR and LCN2-Flag/LCN2 co-immunoprecipitated in HEK293T cells, confirming their interaction with each other (Fig. 4L). This interaction was also observed between the endogenous EGFR and LCN2 in S-CM-treated LOVO cells (Fig. S4E).

**Figure 4.**
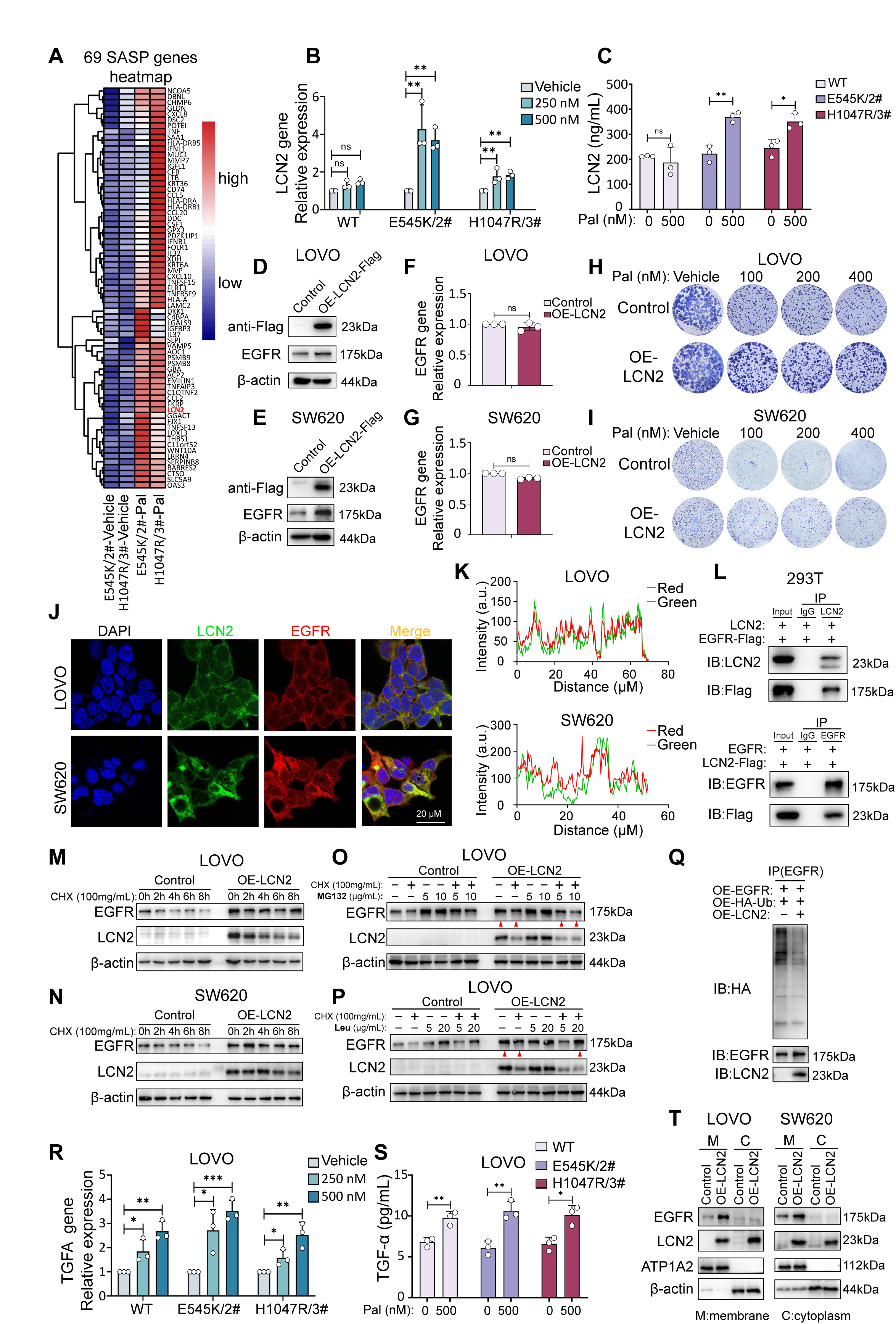
The SASP factor LCN2 enhances the expression of EGFR by suppressing ubiquitination and lysosomal degradation pathways. **A,** The heatmap shows the expression of 69 key SASP genes in PIK3CA mutant cell lines. **B,** mRNA expression of LCN2 genes from LOVO-PIK3CA WT and MUT cell lines treated with palbociclib. **C,** ELISA to measure the secretion of LCN2 protein (ng/mL) in LOVO-PIK3CA WT and MUT cell lines treated with palbociclib. **D-E,** Immunoblot for EGFR expression in LOVO and SW620 cells after overexpression of LCN2 gene. **F-G,** mRNA expression of EGFR genes in LOVO and SW620 cells after overexpression of LCN2 gene. **H-I,** Images showing the colony formation of LOVO and SW620 cells after overexpression of LCN2 gene. **J,** Immunofluorescence images of EGFR (red) and LCN2 (green) LOVO and SW620 cells after overexpression of EGFR and LCN2 genes (Scale bar: 20 μm). **K,** Plot profile analysis of colocalization between EGFR and LCN2 proteins. **L,** Co-IP immunoblot of EGFR-Flag and LCN2-Flag in transfected HEK293T cells. **M-N,** Immunoblot for EGFR expression with treatment CHX (100 mg/mL) in LOVO and SW620 cell lines after overexpression of LCN2 gene and control. **O,** Immunoblot for EGFR expression with treatment CHX (100 mg/mL) and MG132 in LOVO cell lines after overexpression of LCN2 gene and control. **P,** Immunoblot for EGFR expression with treatment CHX (100 mg/mL) and Leu in LOVO cell lines after overexpression of LCN2 gene and control. **Q,** Immunoblot analysis of changes in EGFR ubiquitination under overexpression of EGFR, HA-Ub and LCN2 genes. **R,** mRNA expression of TGFA genes from LOVO-PIK3CA WT and MUT cell lines treated with palbociclib. **S,** ELISA to measure the secretion of TGF-α protein (pg/mL) in LOVO-PIK3CA WT and MUT cell lines treated with palbociclib. **T,** Immunoblot for EGFR expression in the cell membrane and cytoplasm of LOVO and SW620 cells after overexpression of LCN2. ATP1A2 protein was used as a membrane protein positive control. Mean ± SD; *, *P* < 0.05, **, *P* < 0.01; and ***, *P* < 0.001.

To investigate how LCN2 influenced the expression of EGFR, we inhibited protein synthesis using cycloheximide (CHX) and monitored the change in EGFR level in CRC cells. Results showed that LCN2 overexpression significantly slowed EGFR protein degradation in LOVO and SW620 cells (Fig. 4M and N). In contrast, inhibition of overall RNA synthesis by actinomycin D (Act D) failed to prevent the degradation of EGFR in LCN2-overexpressing LOVO cells, suggesting LCN2 upregulates EGFR not via its transcription (Fig. S4F). Next, we treated LOVO cells with a proteasomal inhibitor MG132 or a lysosomal inhibitor leupeptin (Leu), with or without CHX. Results showed that neither 5 nor 10 µg/mL MG132 prevented EGFR degradation (Fig. 4O). In contrast, 20 µg/mL leupeptin completely halted EGFR degradation in LCN2-overexpressing LOVO cells (Fig. 4P), indicating that LCN2 primarily protects EGFR from degradation via the lysosomal pathway. Because ubiquitination regulates protein trafficking and targets cargo proteins for lysosomal degradation ^26^, we examined whether LCN2 affects EGFR ubiquitination and found that LCN2 overexpression did reduce EGFR ubiquitination significantly in 293T cells.

Besides LCN2, TGF-α was critical for EGFR recycling to the plasma membrane and its sustained activation ^25^. We checked the expression and secretion of TGF-α expression and found both levels were significantly increased after palbociclib treatment (Fig. 4R and S). Accordingly, more EGFR was recycled to the plasma membrane in LCN2-overexpressing CRC cells than control in the presence of TGF-α (Fig. 4T), suggesting that LCN2 ultimately enhances the intracellular recycling and membrane localization of EGFR.

### 5. Identification of erlotinib as a palbociclib-synergizer to overcome the palbociclib resistance in PIK3CA mutant CRC cells

To overcome the palbociclib resistance, we utilized a drug screening workflow for rapid identification of drug-drug synergism to identify effective combination therapies for PIK3CA mutant CRC ^27^ (Fig. 5A). In brief, the anti-proliferative activity of 133 clinically relevant small molecule targeted inhibitors and their combined effect with palbociclib in DLD1 cells were assessed, and the Sensitivity Index (SI) of each agent was calculated. Figure 5B shows that 17 agents had an SI above 0, and erlotinib, a selective EGFR inhibitor, achieved the highest SI value, highlighting EGFR’s role in palbociclib resistance (Fig. 5B). To confirm the synergistic interaction between palbociclib and erlotinib, we determined the high single agent (HSA) and combination index (CI) values as readouts for synergistic inhibition in CRC cells carrying intrinsic or exogenous PIK3CA mutation. As a result, most of the HSA values >0 in the concentration combination matrices (Fig. 5C and Fig. S5A) and CI values <1 in the Fa-CI plot in treatment (Fig. 5D and S5B), indicating synergistic interactions occurred in most of the palbociclib-erlotinib combinations at different concentrations. Meanwhile, the addition of erlotinib sensitized the previously resistant DLD1, HCT15, and MUT cells to palbociclib, significantly suppressing their proliferation (Fig. 5E and Fig. S5C) and colony formation (Fig. 5F and Fig. S5D).

**Figure 5.**
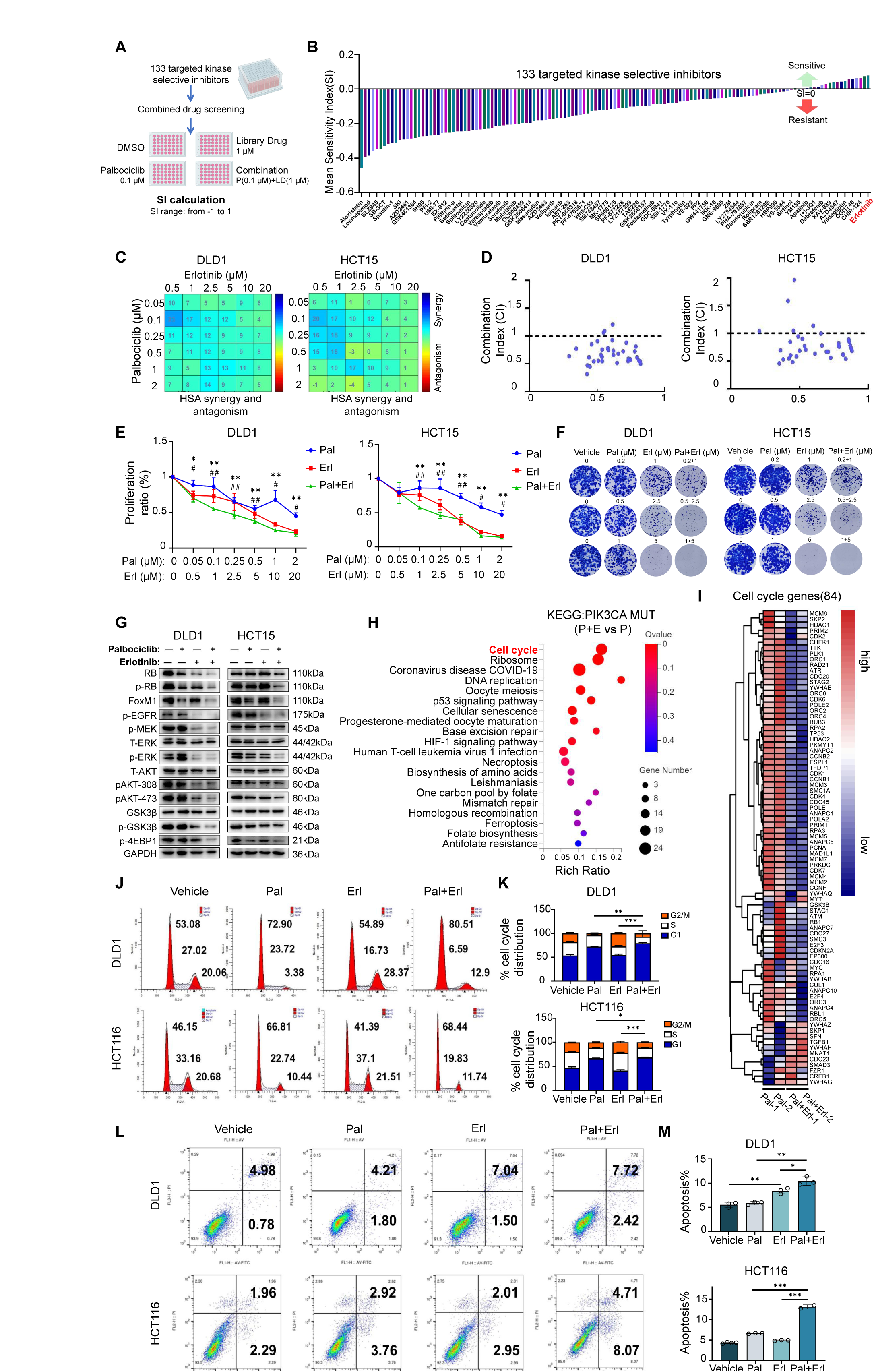
The screening of a drug library for the combination of erlotinib and palbociclib reveals a significant synergistic anti-tumor effect. **A,** Schematic diagram of drug library screening. **B,** Synergy index calculation for 133 tumor-targeting drugs with palbociclib. **C,** HSA synergy matrices showing the interaction between palbociclib and erlotinib in DLD1 and HCT15 cell lines (HSA > 0 indicate synergistic effects). **D,** CI plots showing the sturdy synergistic effect of palbociclib and erlotinib in DLD1 and HCT15 cell lines (CI < 1 indicates a synergistic effects). **E,** Growth inhibition curves of DLD1 and HCT15 cell lines treated with fixed concentration-combinations of palbociclib and erlotinib. **F,** Images showing the colony formation of DLD1 and HCT15 cell lines treated with palbociclib and/or erlotinib. **G,** Immunoblot for EGFR and p-RB downstream protein expression in DLD1 and HCT15 cells following treatment with palbociclib (20 nM), erlotinib (2 μM), or their combination. **H,** KEGG analysis of RNA-seq data comparing palbociclib + erlotinib to palbociclib in LOVO-PIK3CA MUT cells. **I,** Heatmap depicting the expression of 84 cell cycle genes following PE treatment. **J,** Cell cycle analysis of DLD1 and HCT15 cell lines upon treatment with palbociclib or erlotinib alone or in combination. **K,** The quantification of cell fractions in G1, S and G2/M phases in **J**. **L,** Apoptosis analysis of DLD1 and HCT15 cell lines upon treatment of palbociclib and/or erlotinib. Quantification of apoptotic cells in (**M**) by image (**L**). Mean ± SD; *, *P* < 0.05, **, *P* < 0.01; and ***, *P* < 0.001.

To elucidate the underlying mechanism of the palbociclib-erlotinib synergy, we examined the signaling pathways and transcriptome alterations in PIK3CA mutant cells after treatment. Western blotting showed that erlotinib intensified the inhibition of RB and p-RB by palbociclib in CRC cells. Additionally, the combination of palbociclib and erlotinib (PE-combination) further reduced the levels of EGFR and CDK4/6 downstream targets compared with single-agent treatment (Fig. 5G). KEGG analysis of MUT cells treated with palbociclib *vs.* PE-combination showed that the Cell Cycle was the top enriched pathway (Fig. 5H), and 84 representative cell cycle genes were mostly downregulated by PE-combination treatment (Fig. 5I). To investigate the impact of PE-combination on cell cycle arrest and apoptosis, PI/Annexin V-FITC-stained DLD1, HCT15, and MUT cells were analyzed by flow cytometry. Results showed that PE-combination treatment generated a significantly higher proportion of G1-arrested (Fig. 5J-K and Fig. S5E-F) and apoptotic cells (Fig. 5L-M and Fig. S5G-H) than palbociclib-treatment alone.

Cetuximab, an FDA-approved monoclonal antibody targeting EGFR, is used for metastatic or KRAS wild-type CRC, and we explored its potential synergy with palbociclib in treating PIK3CA mutant CRC cells ^28, 29^. The HSA and CI analysis revealed that the interactions between palbociclib and cetuximab were mostly synergistic (Fig. S6A and B) but the antiproliferative activity of cetuximab alone (Fig. S6C) or in combination with palbociclib (Fig. S6D) was only moderate as the half inhibition of the CRC cells’ proliferation was hardly achieved even at a relatively high concentrations (e.g. cetuximab:50 µg/mL; palbociclib:1 µM) (Fig S6E). Since the palbociclib and cetuximab combination was less effective than the PE-combination, cetuximab was not selected for the follow-up efficacy study.

### 6. Combination therapy of palbociclib and erlotinib exhibited significant efficacy in PIK3CA mutant/WT CRC CDX and PDX models

To evaluate the efficacy of the PE-combination therapy *in vivo*, we established PIK3CA mutant CRC PDX models in zebrafish or mice. The tumor tissue transplanted into the 48 hpf Tg (flk1:EGFP) zebrafish embryos or 6-8 weeks immunodeficient NNG mice originated from a 47-year male patient (Ca-2021-6-16, abbr. 616#) diagnosed with papillary adenomas of the rectum at June 2021 and carried a H1047L (gain-of-function) mutation in her PIK3CA gene (Fig. S7A). For the PDX zebrafish model, the PE-combination treatment significantly inhibited the growth of PIK3CA mutant CRC xenograft while neither palbociclib nor erlotinib alone exhibited statistically meaningful difference relative to the control (Fig. 6A and B). For the PDX mouse model, palbociclib (p.o. 25 mg/kg; 3 weeks) alone exerted modest inhibitory effect on 616# tumor growth after two-week treatment (Fig. 6C), but it failed to translate into a meaningful prolongation in overall survival (Fig. 6D). In contrast, the PE-combination treatment (palbociclib: p.o. 25 mg/kg; erlotinib: p.o. 25 mg/kg; 3 weeks) successfully controlled the PIK3CA mutant tumor growth, caused remarkable reduction of tumor volume (Fig. 6C and G), and resulted into markedly prolonged survival compared with both the vehicle-treated and palbociclib-treated PDX mice (Fig. 6D). In addition, efficacy study using a PDO model with the tumor tissue from the same patient confirmed the effectiveness of the PE-combination (Fig. S7B). Next, β-galactosidase and immunohistochemical staining on the tumor tissue from PDX mouse models confirmed that palbociclib treatment promoted the marked increase of β-galactosidase, LCN2, and EGFR (Fig. 6E and F). However, PE-combination treatment reduced the number of β-galactosidase-positive senescent cells, putatively due to its activity to promote cellular apoptosis. Again, the colocalization of EGFR and LCN2 in the PDX tumor cells was observed by immunofluorescence staining of EGFR LCN2, confirming the direct binding of the two proteins (Fig. 6H).

**Figure 6.**
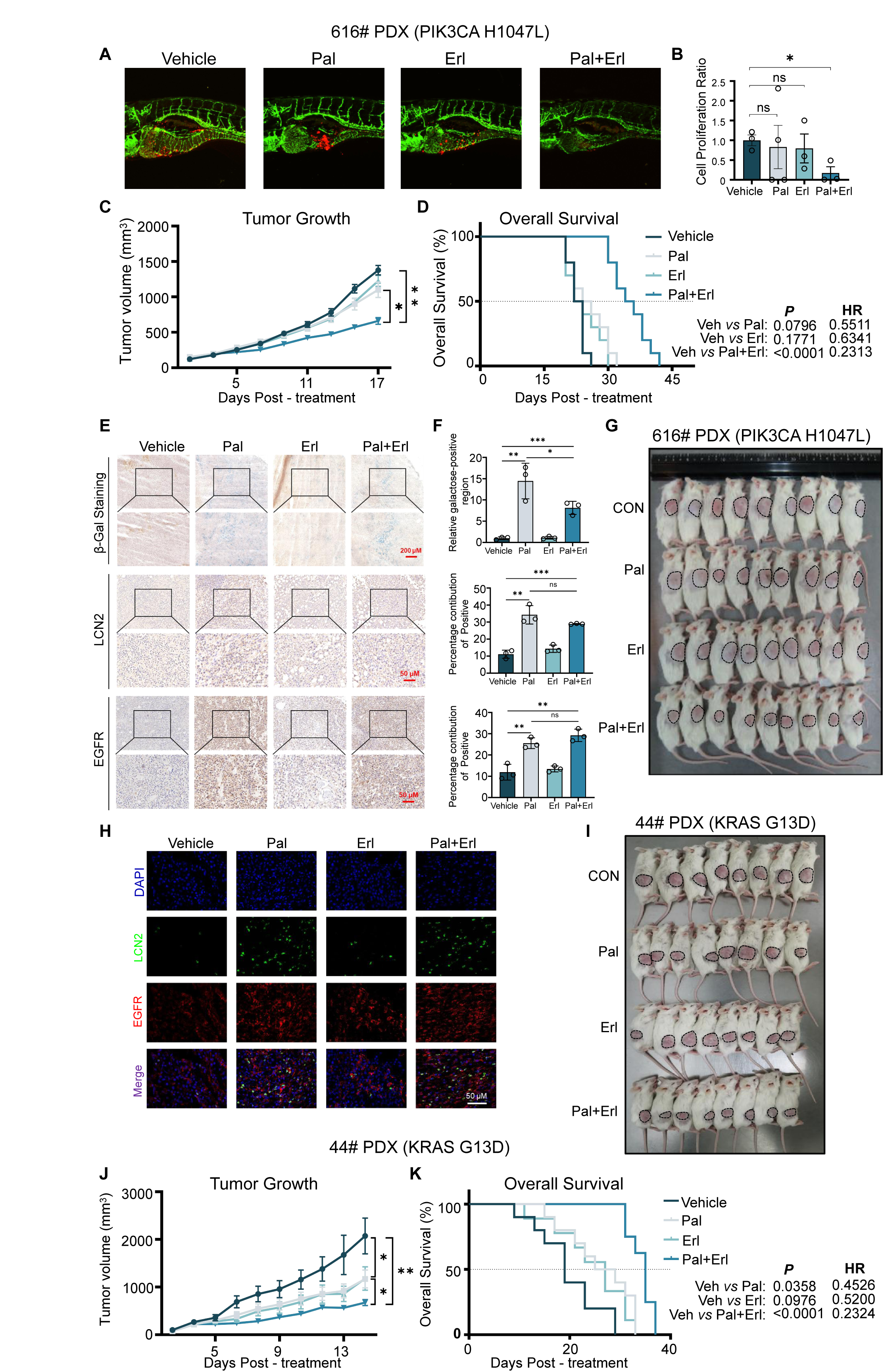
PE-combination exhibited significant efficacy in PDX models with PIK3CA mutation. **A,** Images showing PDX cells with the PIK3CA H1047L mutation treated with palbociclib, erlotinib, or both in zebrafish. Quantification from image (**A**) is shown in **B**. **C,** Tumor volume curves of PDX (616#) after palbociclib and/or erlotinib treatment. **D,** Kaplan-Meier curves depicting OS for PDX (616#) mice treated with palbociclib and/or erlotinib. **E,** PDX (616 #) tumor tissue from mice treated with palbociclib and/or erlotinib, stained for β-Gal, LCN2, and EGFR (Scale bars: 200 µm, 50 µm). Quantification from image (**E**) is shown in **F**. **G,** Representative images of PDX (616#) mice upon treatment of palbociclib and/or erlotinib. **H,** Images showing immunofluorescent staining of PDX (616#) tissues for LCN2 (green) and EGFR (red) proteins after treatment with palbociclib and/or erlotinib (scale bar: 50 µm). **I,** Representative images of PDX (44 #) mice upon treatment of palbociclib and/or erlotinib. **J,** Tumor volume curves of PDX (44#) after palbociclib and/or erlotinib treatment. **K,** Kaplan-Meier curves depicting OS for PDX (44#) mice treated with palbociclib and/or erlotinib. Mean ± SD; Mean ± SEM; *, *P* < 0.05, **, *P* < 0.01; and ***, *P* < 0.001.

**Figure 7.**
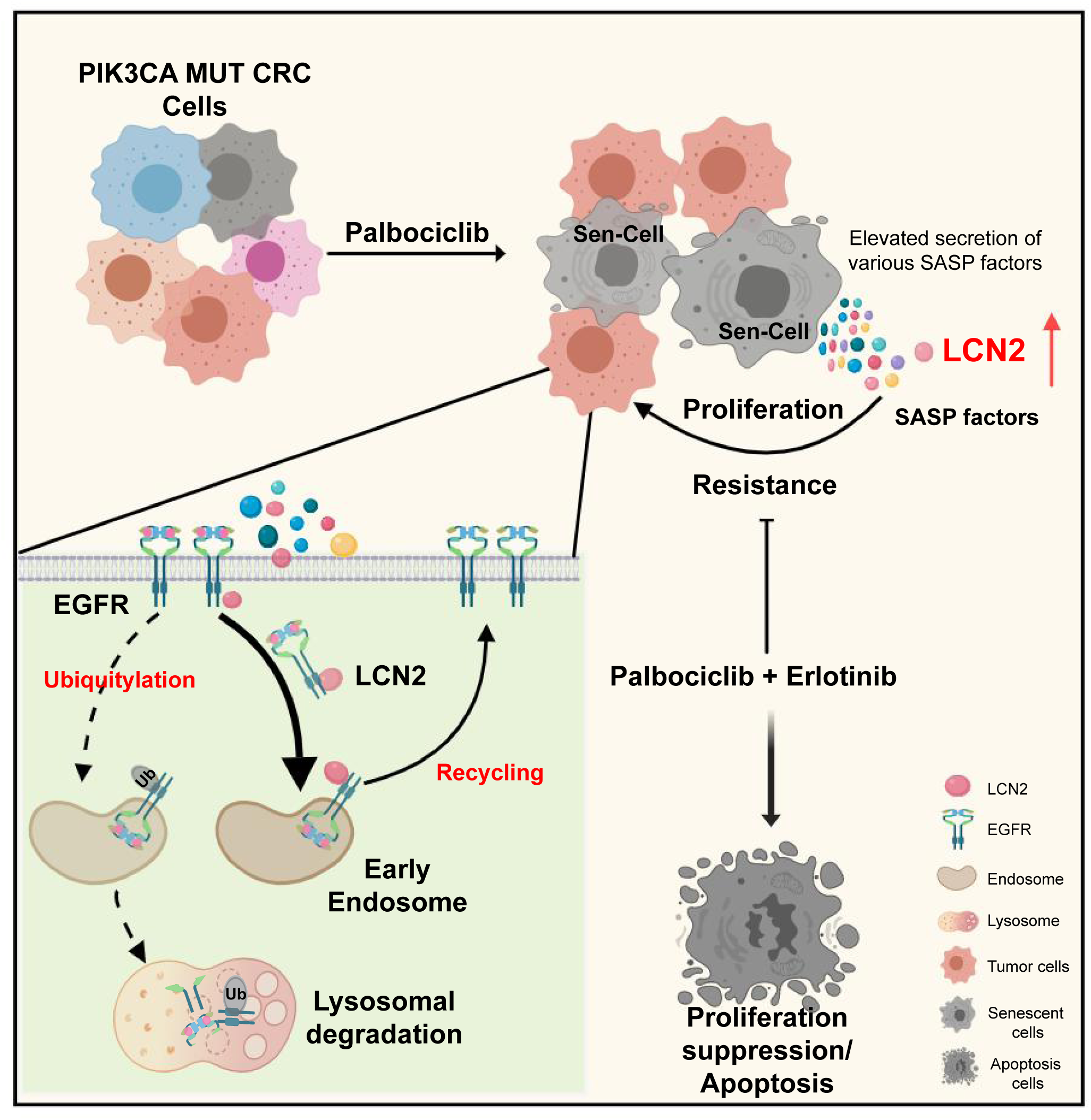
Diagram illustrating the research mechanism. The study’s mechanistic diagram shows that PIK3CA mutations alter the senescence phenotype and SASP gene expression induced by palbociclib, leading to increased EGFR protein expression and enhanced colorectal cancer resistance to the drug. The SASP factor LCN2 is key in upregulating EGFR by inhibiting its ubiquitination and reducing lysosomal degradation. Therapeutically, combining erlotinib with palbociclib could be beneficial.

As the PIK3CA mutant DLD1 cells were refractory to palbociclib *in vitro*, we also evaluated the *in vivo* efficacy of palbociclib and the PE-combination treatment in a DLD1 xenograft model. Again, the single-agent treatment of palbociclib failed to effectively inhibit the tumor growth and prolong the survival of treated mice, whereas the PE-combination treatment resulted in significant tumor suppression and survival prolongation (Fig. S7C-E).

Lastly, we wanted to know whether the PE-combination therapy was also effective for PIK3CA wild-type CRC. To investigate it, tumor tissue derived from a tubulopapillary rectal adenocarcinoma from a 26-year female patient (Ca-18-4-4, abbr. 44#) was transplanted into immunodeficient NNG mice. Sanger sequencing detected a WT PIK3CA gene together with a KRAS G13D mutation in this patient (Fig. S7A). In this PIK3CA WT; KRAS G13D mutant CRC PDX model, palbociclib alone was sufficient to cause significant tumor suppression and survival improvement, but the outcome from the PE-combination group was even more prominent than palbociclib alone (Fig. 6I-K). In addition, H&E and Ki-67 staining of tumor tissues from the DLD1 xenograft, 616#, and 44# models consistently showed that the PE-combination was the most effective of all treatments in restricting the tumor growth (Fig. S7F-H). Therefore, PE-combination therapy can also be a potent and effective treatment for PIK3CA WT CRC. Notably, all mice that received the clinically relevant dose of the PE-combination (palbociclib and erlotinib, each 25 mg/kg orally) showed no severe toxicity or side effects, indicating it is a safe treatment (Fig. S7I).

## DISCUSSION

In the present study, we demonstrated that palbociclib could cause divergent responses in CRC cells depending on their distinct genetic backgrounds. Compared with PIK3CA WT cells, palbociclib treatment led to a strong SASP and altered secretion of SASPF in CRC cells harboring PIK3CA mutation (e.g. E545K and H1047R), and this phenotype was identified as a major contributor to the palbociclib resistance. Mechanistically, the excessively secreted SASPF LCN2 blocked the lysosome-mediated degradation of EGFR in the surrounding non-senescent CRC cells, and the overexpressed EGFR rendered upregulation of multiple palbociclib-resistance genes in PIK3CA mutant CRC cells. Importantly, the SASP-induced resistance could be overcome by concurrent administration of erlotinib and palbociclib. Eventually, this combination therapy significantly suppressed the proliferation and induced apoptosis in both PIK3CA mutant and wt CRC cells, exhibiting potent efficacy and excellent safety in PDO/PDX/CDX models (Fig. 6 and S7).

It is noteworthy that two early studies also reported that the efficacy of palbociclib could be compromised by the PIK3CA mutation in oral squamous cell carcinoma and melanoma ^30, 31^. Although the authors proposed different mechanisms, they probably observed a similar phenomenon as we observed. Tumor cells with OIS have adapted to a pre-senescent state, so when they confronted exogenous stress induced by drug(s) with TIS activity, a survival signal could be triggered and transmitted via the secretion of SASPF to counterbalance the stress. From the perspective of tumor evolution, this change appears to be an adaptive mechanism by which the stress of cell cycle arrest and cellular senescence on the whole tumor cell community can be greatly alleviated and tumor cells survive.

Just as the patients with triple-negative breast cancers could not benefit from the palbociclib, it would be reasonable that only a subgroup of CRC patients is suitable for the palbociclib treatment as well. Therefore, our study highlights the importance of identifying the potential treatment-beneficial/refractory subgroup(s) when a targeted therapeutic agent is involved. Finally, we recently initiated a project aiming to develop a CDK4/6 and EGFR multi-targeting inhibitor using a pharmacophore fusion strategy, and have identified several candidate compounds with promising activities. It is anticipated that these efforts together will eventually translate into effective therapeutic solutions that benefit CRC patients even if they are in a treatment-refractory subgroup.

### Data Availability

The RNA-seq data have been deposited in the Gene Expression Omnibus (GEO) and are publicly accessible as of the publication date.

### Authors’ Disclosures

The authors declare no competing interests.

## Supporting information

Palbociclib resistance genes from literature

---

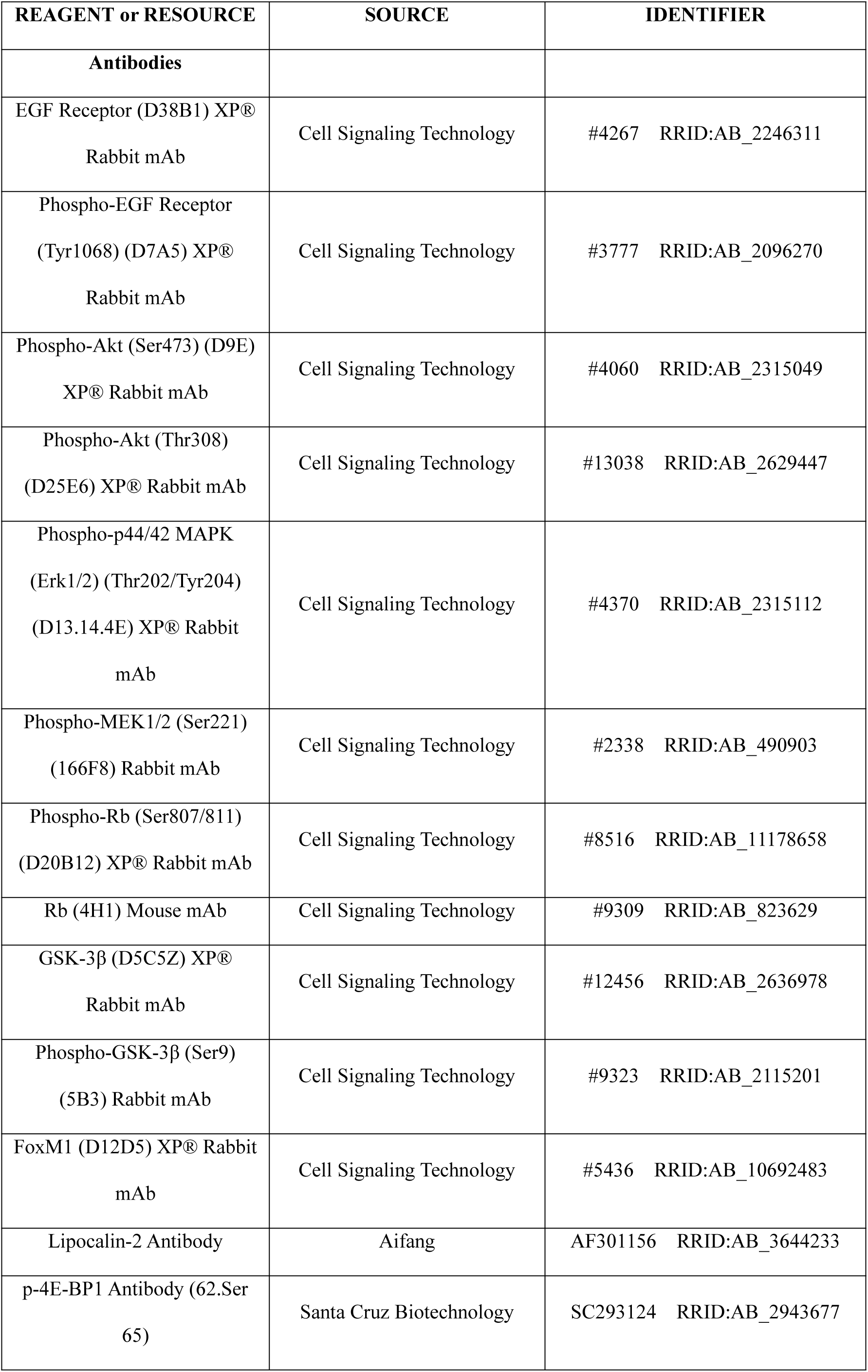
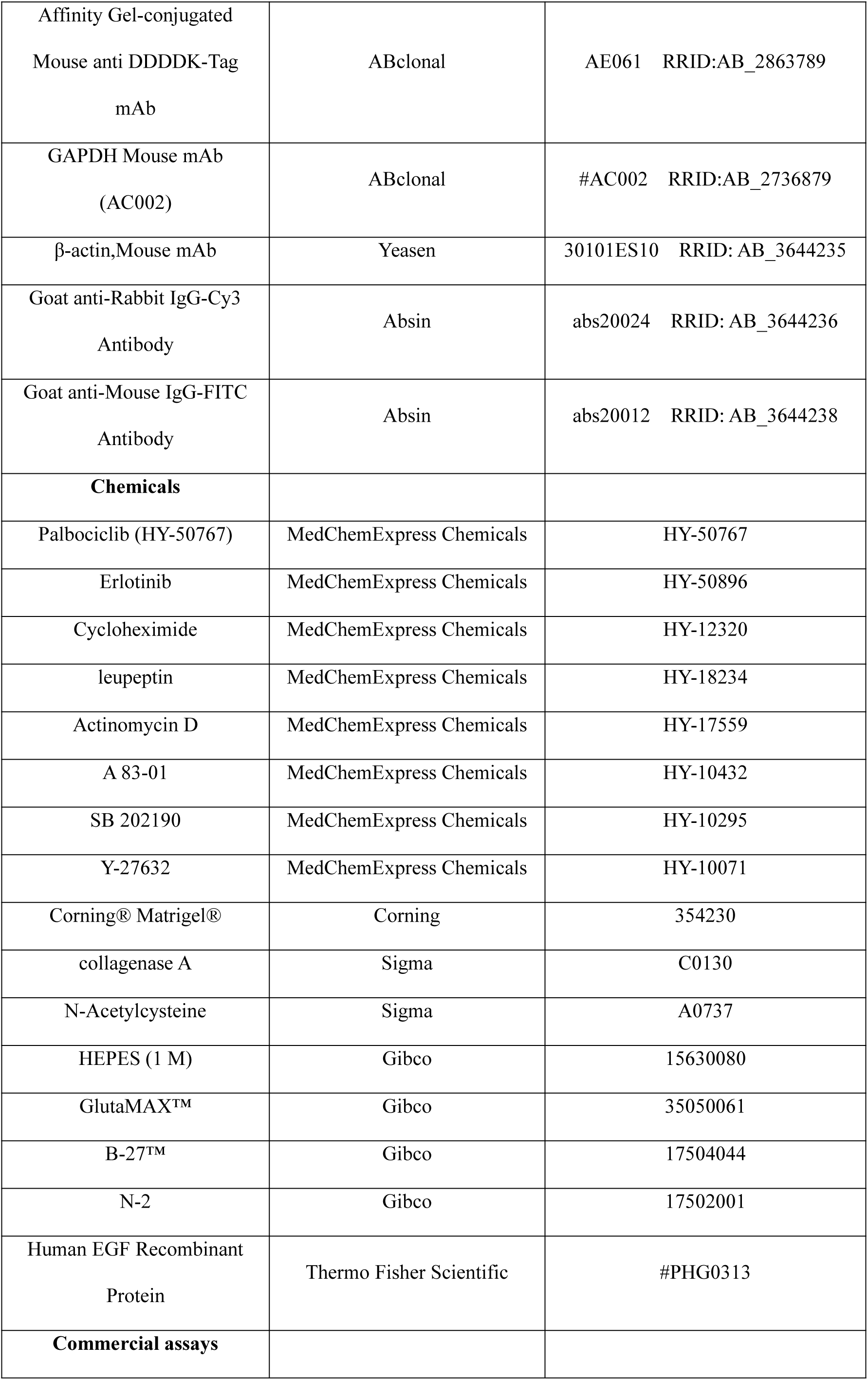
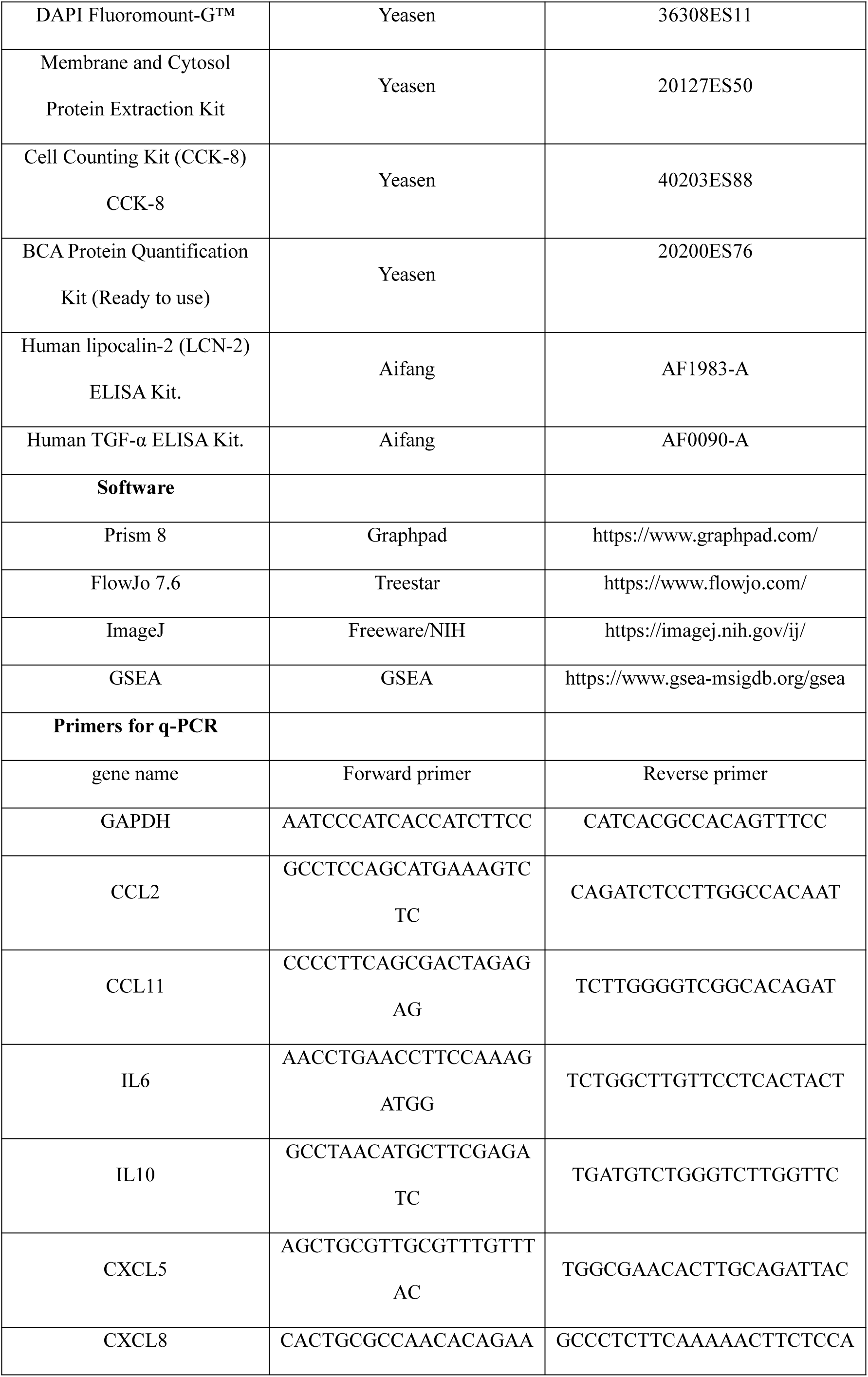
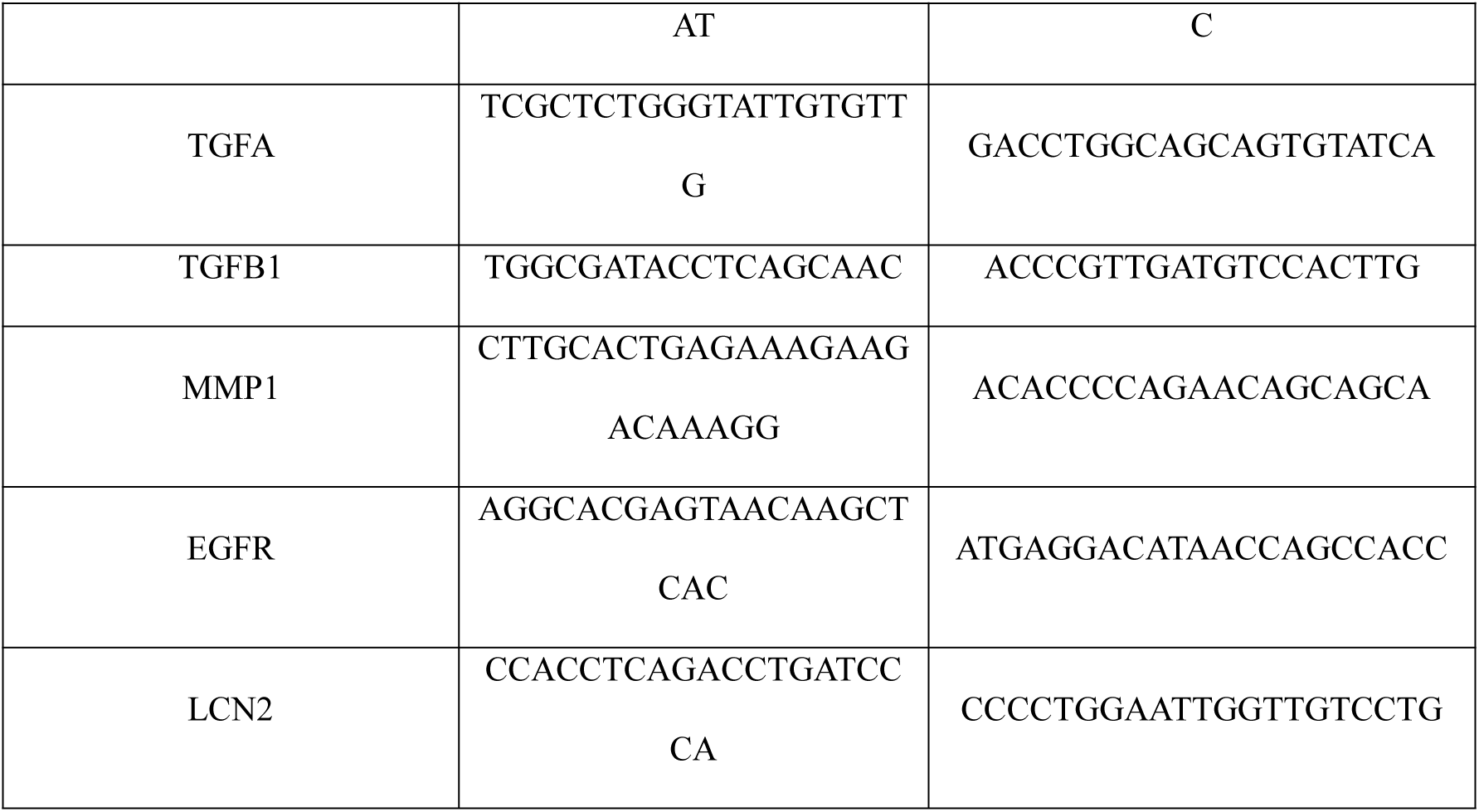
REAGENT or RESOURCE.

## METHODS

### Cell lines

Human colorectal cancer (CRC) cell lines, including SW620, HCT116, HCT15, DLD1, and LOVO, were procured from the Cell Bank of the Chinese Academy of Sciences, located in Shanghai, China. Authentication of these cell lines was conducted by Biowing Biotech, also based in Shanghai, China. The SW620, HCT116, and LOVO cell lines were cultured in Dulbecco’s Modified Eagle Medium (DMEM; Gibco, CA, USA), enriched with 10% fetal bovine serum (FBS; TransGen, Beijing, China), and supplemented with 100 units/mL penicillin and 100 μg/mL streptomycin (Invitrogen, CA, USA). The remaining cell lines, HCT15 and DLD1, were cultured in RPMI-1640 medium (Gibco, CA, USA) with identical supplements. All cells were cultured in a humidified incubator with 5% carbon dioxide (CO_2_) and 95% air at 37 °C and were routinely checked for mycoplasma by PCR analysis.

### Plasmid construction

A lentiviral vector targeting the PIK3CA gene’s E545K and H1047R mutations was systematically developed. The process began with synthesizing and amplifying the human wild-type PIK3CA expression vector plvx using PCR. Then, plasmids with E545K and H1047R mutations were created via homologous recombination using the CloneExpress Ultra One Step Cloning Kit (Vazyme, Nanjing, China), following the manufacturer’s instructions.

### Lentivirus production and infection

HEK293T cells were plated in 10-cm dishes at 2 × 10^6^ cells per dish and incubated for 24 hours. When the cell density reached 60% – 70%, a mixture containing plvx, pVSVg, and psPAX2 was transfected to each 10-cm dish of HEK293T cells. After 18 hours, the medium was refreshed. The lentivirus-containing supernatant was collected and filtered 48 hours later. Cells were seeded in 12-well plates for 24 hours and then infected with lentivirus by centrifuging at 2000 rpm for 2 hours at 37 °C in the presence of 8 μg/ml polybrene to increase infection efficiency.

### Collection of conditioned media (CM)

LOVO cells were seeded in 10-cm dishes and cultured overnight. They were then treated with 0.5 μM palbociclib and incubated for 3 days. After washing with phosphate-buffered saline (PBS) and incubating in fresh medium for 48 hours, the supernatant was collected, centrifuged at 3000 rpm for 5 minutes, aliquoted, and stored at –80 °C. This preparation was called senescent cell-conditioned medium (S-CM), with DMSO-treated conditioned medium as the control.

### Clonogenic assays

Colorectal cancer cells were cultured in 12-well plates at 500-1000 cells per well, with the medium (drug-treated or fresh) changed every 3-4 days. After 10-14 days, colonies were washed with PBS, fixed with 4% paraformaldehyde for 15 minutes, and stained with Giemsa Stain (KeyGEN, Nanjing, China).

### Proliferation and anti-proliferation assays

Colorectal cancer cells were seeded into 96-well plates (1,000-3,000 cells/well) and cultured for 24 hours. They were then treated with various concentrations of palbociclib and/or erlotinib for 72 hours. After treatment, the medium was replaced with fresh medium containing CCK8 (10 μL/well). After an additional incubation period of 1 to 4 hours, the optical density at 450 nm for each well was measured using a microplate reader.

### RNA extraction and quantitative real-time PCR assay (qRT-PCR)

RNA was extracted with Trizol (Takara, Japan) as per the manufacturer’s guidelines. Then, 1000 ng of RNA was reverse-transcribed into cDNA following Takara’s instructions. qRT-PCR was performed using SYBR Green q-PCR mix (Yeasen, Shanghai, China). Gene mRNA levels were normalized to GAPDH and calculated using the 2^(–ΔΔCt) method, with data analysis done in GraphPad Prism.

### Western blotting

Cells were lysed using RIPA buffer supplemented with protease inhibitors. Protein concentrations were quantified utilizing the BCA Protein Assay Kit (Yeasen, Shanghai, China). Subsequently, equal amounts of protein (10-20 μg) were subjected to SDS-PAGE for electrophoretic separation. Following electrophoresis, proteins were transferred onto a PVDF membrane (Merck Millipore, MA, USA). The membrane was then blocked with 3% bovine serum albumin (BSA) and incubated with appropriate primary and secondary antibodies. Imaging of the membranes was conducted using a Tanon Imager (Tanon, Shanghai, China).

### RNA sequencing

Total RNA was extracted from cells using TRIzol® Reagent as per Invitrogen’s protocol, followed by library construction and RNA sequencing by BGI (Beijing, China). Detailed methods, raw data, and gene expression profiles are available in the GEO database.

### Drug combination screening

A library of 133 target-selective small molecule inhibitors was obtained from Selleck Chemicals. DLD1 cells were plated in 96-well plates at 2000 cells per well, 24 hours before screening. The drug screen involved treating cells with DMSO (0.2 %), palbociclib (0.1 μM), a library agent (1 μM), and a combination of palbociclib and the targeted agent. Sensitivity Index (SI) values were calculated using the formula SI = ECE (Expected Combined Effect) – OCE (Observed Combined Effect), where SI > 0 indicates sensitivity and SI < 0 indicates resistance.

### Mass spectrometry analysis

Cells were lysed in 8 M urea with 50 mM Tris buffer and protease inhibitors, then sonicated. Debris was removed by centrifugation at 20,000 g for 30 minutes at 4 °C. Protein concentration was determined using the Bradford assay. 100 μg of protein was reduced with DTT, alkylated with iodoacetamide, and digested with trypsin. Peptides were desalted with C18 columns and dried. They were separated on a 75 μm × 16 cm C18 column using the Easy-nLC 1200 system. The analysis utilized an Orbitrap Fusion Lumos mass spectrometer (Thermo Fisher) in DIA mode, with MS1 scans at 350-1500 m/z and 120,000 resolution, and MS/MS using HCD at 30,000 resolution. DIA-NN v1.8 processed the DIA files without a library ^39^, referencing the human UniProt database (2023_05) with default settings. Peptide and protein FDR was controlled at 1% using a target-decoy method.

### ELISA assays

Following a 72-hour treatment of LOVO cells with palbociclib, the culture medium was refreshed. Subsequently, after an additional 48-hour period, the concentrations of LCN2 and TGF-α proteins present in the cell supernatant were quantified utilizing ELISA kits (Aifang, Wuhan, China) following the manufacturer’s protocol.

### Co-immunoprecipitation experiments

Immunoprecipitation assays were performed on 293T and LOVO cell lines. Cells were co-transfected with 5 µg each of LCN2-Flag and EGFR-Flag plasmids using polyethyleneimine. After 48 hours, cells were lysed, and the supernatant was used for overnight immunoprecipitation at 4 °C with EGFR and LCN2 antibodies, using rabbit IgG as a negative control. The next day, magnetic beads were added to the mixture and incubated for 2 hours at 4 °C. After incubation, the beads were washed, and proteins were eluted, denatured, and analyzed by SDS-PAGE electrophoresis.

### Cell immunofluorescence

Immunofluorescence was performed on 293T cells co-transfected with LCN2-Flag and EGFR-Flag plasmids using polyethyleneimine. After 48 hours, cells were washed with PBS, fixed with 4% paraformaldehyde for 30 minutes, blocked with 5% goat serum for 1 hour, and incubated overnight with primary antibodies for LCN2 and EGFR. They were then treated for 2 hours at room temperature with fluorescent secondary antibodies: Goat anti-Rabbit IgG-Cy3 and Goat anti-Mouse IgG-FITC. Slides were mounted with a DAPI-containing medium and imaged.

### Membrane protein extraction

LOVO and SW620 cells, stably transfected with the LCN2 gene (1 × 10^6^ cells), were collected, washed with PBS, and scraped from the culture dish. They were centrifuged at 200 × g for 5 minutes at 4 °C to obtain a pellet. Proteins were subsequently extracted from the cell membrane and cytoplasm utilizing the membrane and cytoplasm protein extraction kit (Yeasen, Shanghai, China) in accordance with the manufacturer’s instructions.

### Ubiquitination assays

HEK 293T cells were co-transfected with plasmids for EGFR-Flag, LCN2-Flag, and HA-Ubiquitin to study EGFR ubiquitination. After 48 hours, cells were lysed in a buffer, incubated on ice for 20 minutes, and centrifuged at 12,000 rpm for 15 minutes. The supernatant underwent overnight immunoprecipitation at 4 °C with anti-EGFR antibodies. Afterward, magnetic beads were added, and the mixture was incubated for an additional 2 hours at 4 °C. The Protein A/G Magnetic Beads used for immunoprecipitation (Selleck, USA) were washed, and proteins were eluted and denatured. SDS-PAGE electrophoresis was then performed. Western blot analysis with anti-Flag and anti-HA antibodies was used to detect HA, LCN2, and EGFR in the precipitates.

### β-Gal staining

Intracellular β-galactosidase levels were measured using a Cell Senescence Testing Kit (GenMed Scientifics, MN, USA) according to the manufacturer’s instructions. CRC cells, pre-seeded in 12-well plates, were treated with palbociclib for 72 hours. They were then fixed, washed, and stained for 12-16 hours at 37 °C. After three PBS washes, the cells were observed under a Zeiss microscope.

### Flow Cytometry Analysis of Cell Cycle and Apoptosis

Cells were seeded in 6-well plates at 1 × 10^5^ cells per well and cultured for 24 hours. For cell cycle analysis, they were synchronized by 24-hour serum starvation in serum-free DMEM. Then, cells were treated with palbociclib, erlotinib, or both for 72 hours. Post-treatment, cells were trypsinized into ice-cold PBS, centrifuged, washed with PBS, and fixed in 75% ethanol at –20 °C for 12 hours. A cell cycle analysis kit was used for staining. For apoptosis analysis, cells were treated with varying concentrations of palbociclib or its formulations (palbociclib, erlotinib) for 72 hours. Post-treatment, cells were stained for 20 minutes using the Annexin V-FITC kit as per the manufacturer’s instructions and then collected for flow cytometry analysis.

### Animals

The mice were housed and handled following institutional and local guidelines, with experiments approved by Nanjing Medical University’s Animal Experiment Committee. NOD/ShiLtJGpt-*Prkdc^em26Cd52^Il2rg^em26Cd^*^22^/Gpt (NNG, 3–4 weeks) were sourced from a local center and acclimated for a week before experiments. Transgenic zebrafish embryos (Tg(flk1:EGFP)) were provided by Nanjing HuanTe Biotechnology Co., Ltd.

### Xenograft

For the Cell line-derived xenograft (CDX) model, 1 × 10^6^ LOVO-PIK3CA WT/MUT cells or DLD1 cells were injected into the right flank of each mouse to create a CRC xenograft model. Once tumors averaged 100-120 mm³, mice were divided into groups for experiments. For the patient-derived xenograft (PDX) mouse model study, 4-5 week-old female immunodeficient NNG mice received patient-derived CRC tissue. CRC tissues from two patients were cut into 0.3 cm³ pieces and implanted subcutaneously into the right axillary area of each mouse. For the PDX zebrafish model, 48-hour-old Tg (flk1:EGFP) transgenic zebrafish embryos were injected with 400 tumor cells each to create a transplantation model. These embryos were incubated at 34 °C with a 14-hour light and 10-hour dark cycle for 24 hours. Post-treatment, they were incubated at 37 °C for 72 hours, followed by confocal imaging and proliferation analysis with GraphPad Prism software.

### Drug administration

For the CDX and PDX models, once tumors reached 80-120 mm³, mice were grouped and treated with erlotinib (25 mg/kg in 6% Captisol), palbociclib (25 mg/kg in sodium-lactate buffer, pH 4.0), or both, 5 days a week for 3 weeks. Tumor size (length × width ^2^ × 0.5) and mouse weight were recorded every 2 days, and tumor volume was calculated until it reached 2,000 mm³ or the mice’s health declined significantly. For the PDX zebrafish model, They were then grouped for treatment with DMSO, palbociclib (0.5 μM), erlotinib (1 μM), or a combination of these drugs. Post-treatment, they were incubated at 37 °C for 72 hours, followed by confocal imaging and proliferation analysis with GraphPad Prism software.

### Patient-derived CRC organoids

CRC specimens were collected with informed consent at Jiangsu Cancer Hospital, and organoids were created following Hans Clevers’ protocol ^40^. Tissues were cut into 1 mm³ pieces, digested with collagenase A, centrifuged, and resuspended in PBS. The cells were filtered, centrifuged, and mixed with matrigel at 1 × 10^6^ cells/mL. This mixture was placed as 25 μL drops in 24-well plates and solidified at 37 °C for 10 minutes. Each well was filled with 500 μL of culture medium containing niche factors, including EGF, TGF-β receptor I inhibitor, and p38 MAPK inhibitor. Y-27632 was added for the first two days. After CRC organoids formed, palbociclib, erlotinib, or both were introduced to evaluate their effects.

### Bioinformatic analysis

The BGI Genomics platform, Dr. Tom (https://biosys.bgi.com), is used for KEGG, GO, and GSEA analyses. BioLadder (https://www.bioladder.cn/web/#/pro/cloud) creates volcano plots, and STRING (https://cn.string-db.org) analyzes EGFR protein interactions. Heatmaps are generated in Rstudio using Pretty Heatmaps and Ggplot2, while GSEA software analyzes high-throughput data.

### Statistical analysis

The *in vitro* data in the figures are represented as the mean ± SD and the data of PDX tumor growth are presented as mean ± SEM (standard error of the mean). Statistical significance was calculated by Student’s t-test or ANOVA using GraphPad Prism. The *in vitro* experiments were performed in triplicate and replicated in more than two independent experiments. The statistical significance of differences is indicated in figures by asterisks as follows: *, *P* <0.05; **, *P* <0.01; ***, *P* <0.001.

## REFERENCES

1. Sung H et al. Global Cancer Statistics 2020: GLOBOCAN Estimates of Incidence and Mortality Worldwide for 36 Cancers in 185 Countries. CA Cancer J Clin. 71, 209–249 (2021).

2. Biller LH, Schrag D. Diagnosis and Treatment of Metastatic Colorectal Cancer: A Review. JAMA. 325, 669–685 (2021).

3. Beaver JA et al. FDA Approval: Palbociclib for the Treatment of Postmenopausal Patients with Estrogen Receptor-Positive, HER2-Negative Metastatic Breast Cancer. Clin Cancer Res. 21, 4760–4766 (2015).

4. Sorah JD et al. Phase II Single-Arm Study of Palbociclib and Cetuximab Rechallenge in Patients with KRAS/NRAS/BRAF Wild-Type Colorectal Cancer. Oncologist. 27, 1006–e1930 (2022).

5. Sorah JD et al. Phase II Single-Arm Study of Palbociclib and Cetuximab Rechallenge in Patients with Wild-Type Colorectal Cancer. Oncologist. 27, 1006–E1930 (2022).

6. Matthews HK, Bertoli C, de Bruin RAM. Cell cycle control in cancer. Nat Rev Mol Cell Biol. 23, 74–88 (2022).

7. Goel S, Bergholz JS, Zhao JJ. Targeting CDK4 and CDK6 in cancer. Nat Rev Cancer. 22, 356–372 (2022).

8. Thoma OM, Neurath MF, Waldner MJ. Cyclin-Dependent Kinase Inhibitors and Their Therapeutic Potential in Colorectal Cancer Treatment. Front Pharmacol. 12, 757120 (2021).

9. Goel S, DeCristo MJ, McAllister SS, Zhao JJ. CDK4/6 Inhibition in Cancer: Beyond Cell Cycle Arrest. Trends Cell Biol. 28, 911–925 (2018).

10. Wang B et al. Pharmacological CDK4/6 inhibition reveals a p53-dependent senescent state with restricted toxicity. EMBO J. 41, e108946 (2022).

11. Vijayaraghavan S et al. CDK4/6 and autophagy inhibitors synergistically induce senescence in Rb positive cytoplasmic cyclin E negative cancers. Nat Commun. 8, 15916 (2017).

12. Rossiello F, Jurk D, Passos JF, d’Adda di Fagagna F. Telomere dysfunction in ageing and age-related diseases. Nat Cell Biol. 24, 135–147 (2022).

13. Ou HL et al. Cellular senescence in cancer: from mechanisms to detection. Mol Oncol. 15, 2634–2671 (2021).

14. Guillon J et al. Chemotherapy-induced senescence, an adaptive mechanism driving resistance and tumor heterogeneity. Cell Cycle. 18, 2385–2397 (2019).

15. Zhu H et al. Oncogene-induced senescence: From biology to therapy. Mech Ageing Dev. 187, 111229 (2020).

16. Schmitt CA, Wang BS, Demaria M. Senescence and cancer – role and therapeutic opportunities. Nat Rev Clin Oncol. 19, 619–636 (2022).

17. Faget DV, Ren Q, Stewart SA. Unmasking senescence: context-dependent effects of SASP in cancer. Nat Rev Cancer. 19, 439–453 (2019).

18. Birch J, Gil J. Senescence and the SASP: many therapeutic avenues. Gene Dev. 34, 1565–1576 (2020).

19. Courlet P et al. Population Pharmacokinetics of Palbociclib and Its Correlation with Clinical Efficacy and Safety in Patients with Advanced Breast Cancer. Pharmaceutics. 14, (2022).

20. Tan JX, Finkel T. Lysosomes in senescence and aging. EMBO Rep. 24, e57265 (2023).

21. Guerrero-Navarro L, Jansen-Durr P, Cavinato M. Age-Related Lysosomal Dysfunctions. Cells. 11, (2022).

22. Lasry A, Ben-Neriah Y. Senescence-associated inflammatory responses: aging and cancer perspectives. Trends Immunol. 36, 217–228 (2015).

23. Chen C et al. Intracavity generation of glioma stem cell-specific CAR macrophages primes locoregional immunity for postoperative glioblastoma therapy. Science translational medicine. 14, eabn1128 (2022).

24. Park YH et al. Longitudinal multi-omics study of palbociclib resistance in HR-positive/HER2-negative metastatic breast cancer. Genome Med. 15, (2023).

25. Yammine L et al. Lipocalin-2 Regulates Epidermal Growth Factor Receptor Intracellular Trafficking. Cell Rep. 29, 2067–2077 e2066 (2019).

26. Alwan HAJ, van Leeuwen JEM. UBPY-mediated epidermal growth factor receptor (EGFR) de-ubiquitination promotes EGFR degradation. Journal of Biological Chemistry. 282, 1658–1669 (2007).

27. Chen Z et al. A high-throughput drug combination screen identifies an anti-glioma synergism between TH588 and PI3K inhibitors. Cancer Cell Int. 20, 337 (2020).

28. Goldberg RM. Cetuximab. Nat Rev Drug Discov. **Suppl**, S10–11 (2005).

29. Goldberg RM, Venook AP, Schilsky RL. Cetuximab in the treatment of colorectal cancer. Clin Adv Hematol Oncol. 2, 1–10; quiz 11-12 (2004).

30. Romano G et al. A Preexisting Rare Subpopulation Confers Clinical Resistance to MEK plus CDK4/6 Inhibition in Melanoma and Is Dependent on S6K1 Signaling. Cancer Discovery. 8, 556–567 (2018).

31. Zainal NS et al. Effects of palbociclib on oral squamous cell carcinoma and the role of PIK3CA in conferring resistance. Cancer Biol Med. 16, 264–275 (2019).

