## Supplemental figure1-7 for "Oncogene-induced senescence and senescence-associated secretory phenotype (SASP) determine the efficacy of palbociclib in PIK3CA mutated colorectal cancer"

Fig. S1

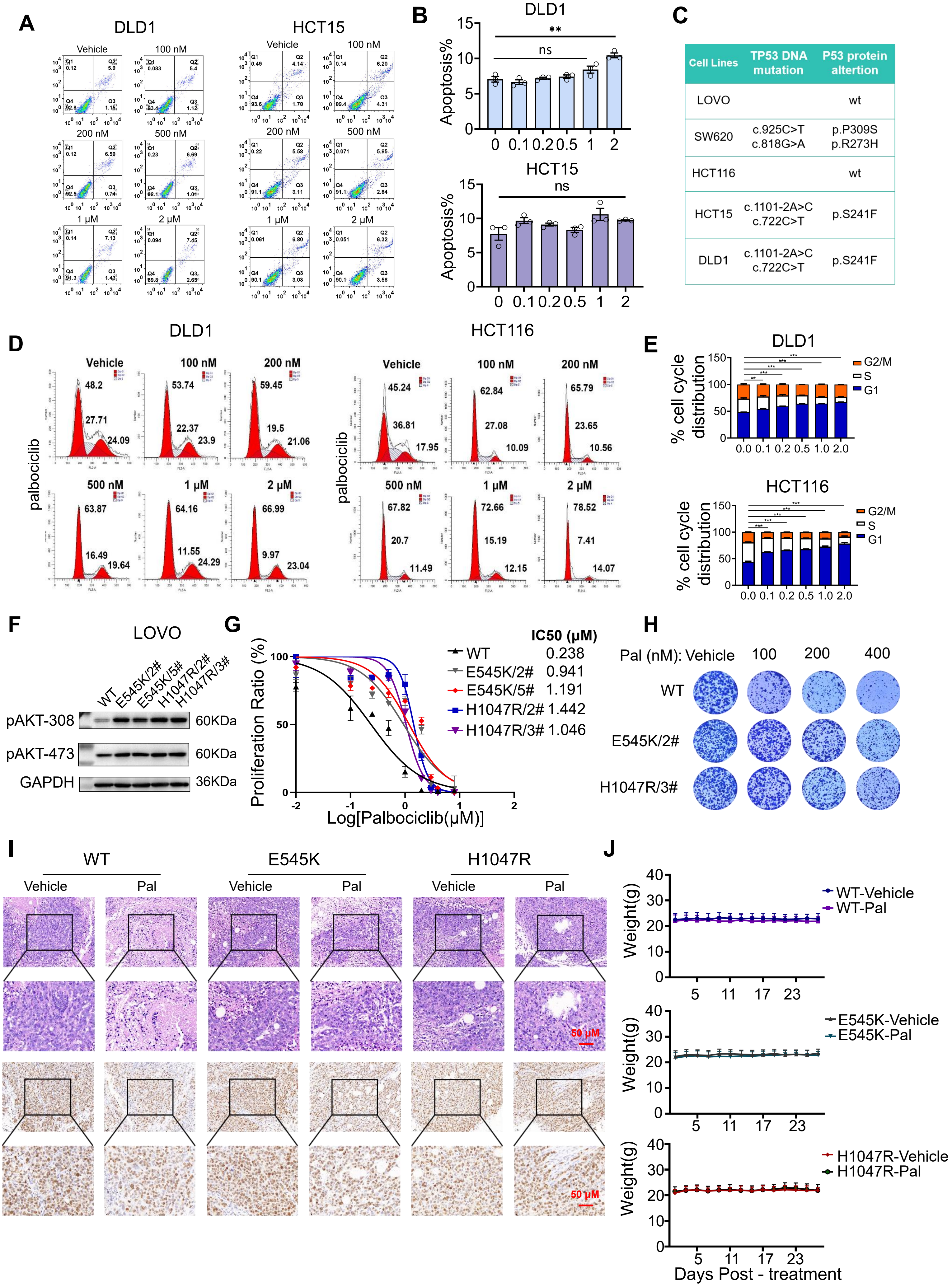
A
DLD1
HCT15
Vehicle
100 nM
200 nM
500 nM
1 μM
2 μM
B
DLD1
HCT15
Apoptosis%
ns
**
C
Cell Lines | TP53 DNA mutation | P53 protein altertion
LOVO | | wt
SW620 | c.925C>T c.818G>A | p.P309S p.R273H
HCT116 | | wt
HCT15 | c.1101-2A>C c.722C>T | p.S241F
DLD1 | c.1101-2A>C c.722C>T | p.S241F
D
DLD1
HCT116
palbociclib
E
% cell cycle distribution
G2/M
S
G1
F
LOVO
WT
E545K/2#
E545K/5#
H1047R/2#
H1047R/3#
pAKT-308
pAKT-473
GAPDH
60KDa
60KDa
36KDa
G
IC50 (μM)
WT 0.238
E545K/2# 0.941
E545K/5# 1.191
H1047R/2# 1.442
H1047R/3# 1.046
Proliferation Ratio (%)
Log[Palbociclib(μM)]
H
Pal (nM): Vehicle 100 200 400
WT
E545K/2#
H1047R/3#
I
WT
E545K
H1047R
Vehicle
Pal
H&E
Ki67
50 μM
J
WT-Vehicle
WT-Pal
E545K-Vehicle
E545K-Pal
H1047R-Vehicle
H1047R-Pal
Weight(g)
Days Post - treatment

**Figure S1. Palbociclib antitumor activity and PIK3CA mutations promote palbociclib resistance in colorectal cancer.**

(**A**) Table of TP53 gene mutations in five CRC cell lines from the CCLE database. (**B**) Apoptosis assessment of DLD1 and HCT15 cells treated with palbociclib, with quantification shown in image **C**. (**D**) Cell cycle analysis of DLD1 and HCT116 cell lines upon treatment with palbociclib. **E,** The quantification of cell fractions in G1, S and G2/M phases in **D**. (**F**) Immunoblot for phosphorylation levels at AKT sites Thr308 and Ser473 in LOVO cell lines stably expressing E545K (2# and 5#) and H1047R (2# and 3#) mutations of the PIK3CA gene. (**G**) Growth inhibition curves and IC50 values for LOVO-PIK3CA MUT cell lines under palbociclib treatment. (**H**) Images showing the colony formation of LOVO-PIK3CA MUT cell lines with palbociclib treatment. (**I**) Tumor tissue stained with H&E and Ki-67 of LOVO-PIK3CA WT and MUT cells mouse CDX upon treatment of palbociclib (scale bar: 50 μm). (**J**) Body weight curves in LOVO-PIK3CA WT and MUT cell mouse CDX following palbociclib treatment. Mean ± SD; *, $P < 0.05$, **, $P < 0.01$; and ***, $P < 0.001$

Fig. S2

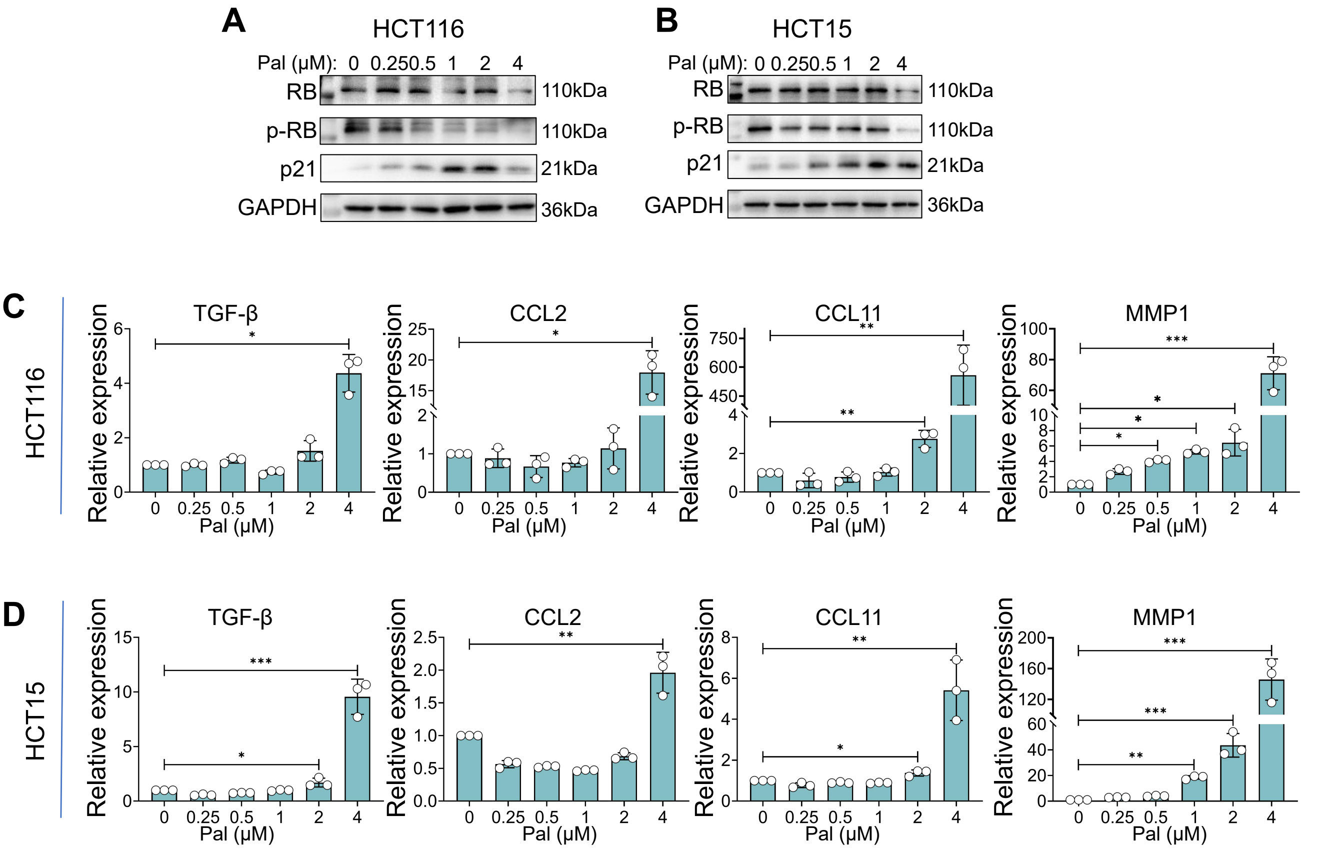

A
HCT116
Pal (μM): 0 0.25 0.5 1 2 4
RB 110kDa
p-RB 110kDa
p21 21kDa
GAPDH 36kDa
B
HCT15
Pal (μM): 0 0.25 0.5 1 2 4
RB 110kDa
p-RB 110kDa
p21 21kDa
GAPDH 36kDa
C
HCT116
TGF-β
CCL2
CCL11
MMP1
Relative expression
Pal (μM)
D
HCT15
TGF-β
CCL2
CCL11
MMP1
Relative expression
Pal (μM)

**Figure S2. Palbociclib induces senescence and SASP gene expression in cells with natural PIK3CA mutation.**

(**A**) Immunoblotting for RB, p-RB, and p21 was conducted on HCT116 and HCT15 cells treated with palbociclib. (**B**) mRNA expression of SASP genes in HCT116 and HCT15 cells treated with palbociclib. Mean ± SD; *, $P < 0.05$, **, $P < 0.01$; and ***, $P < 0.001$

Fig. S3

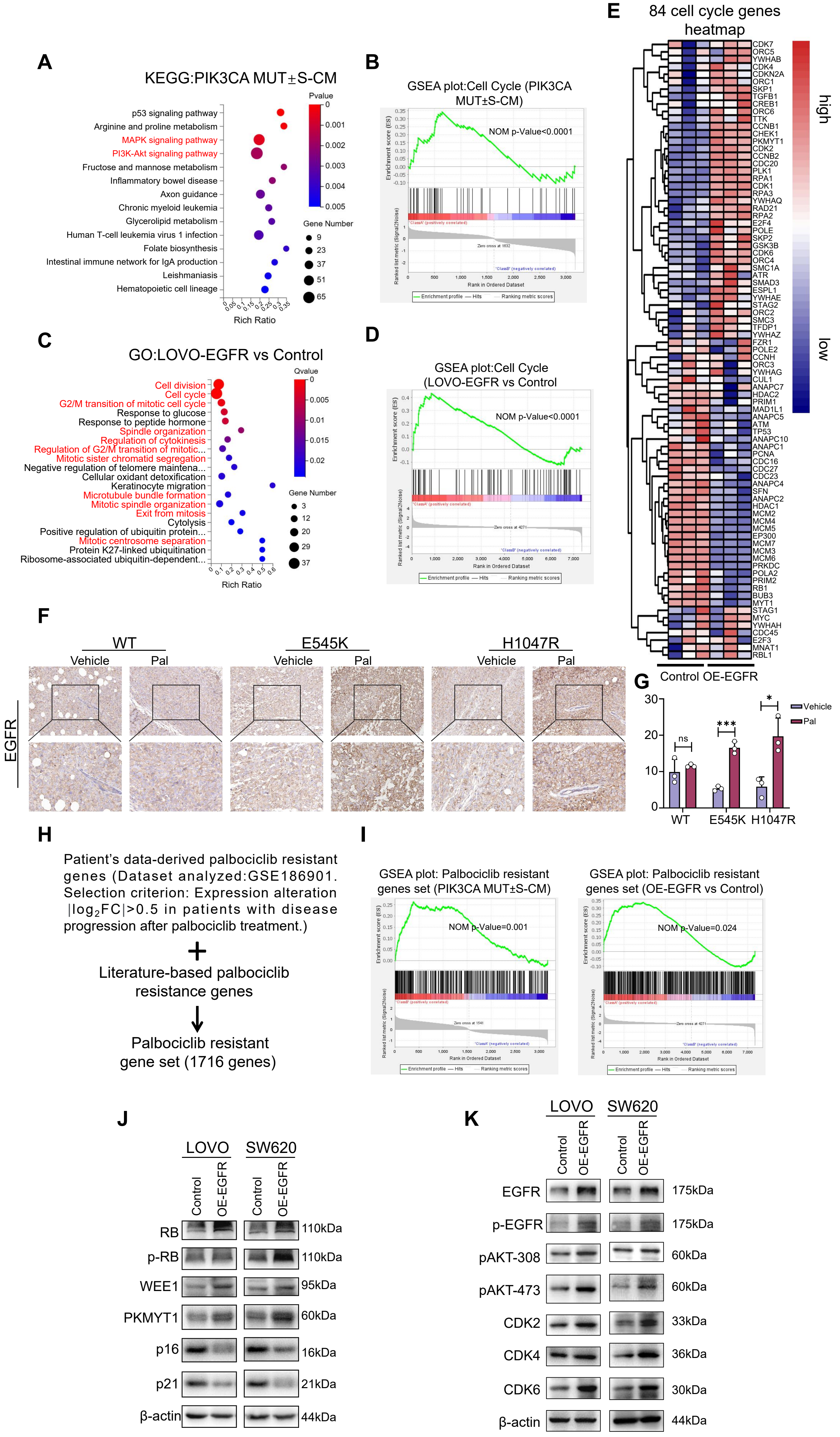
A
KEGG:PIK3CA MUT±S-CM
p53 signaling pathway
Arginine and proline metabolism
MAPK signaling pathway
PI3K-Akt signaling pathway
Fructose and mannose metabolism
Inflammatory bowel disease
Axon guidance
Chronic myeloid leukemia
Glycerolipid metabolism
Human T-cell leukemia virus 1 infection
Folate biosynthesis
Intestinal immune network for IgA production
Leishmaniasis
Hematopoietic cell lineage
Rich Ratio
Pvalue
Gene Number
B
GSEA plot:Cell Cycle (PIK3CA MUT±S-CM)
NOM p-Value<0.0001
C
GO:LOVO-EGFR vs Control
Cell division
Cell cycle
G2/M transition of mitotic cell cycle
Response to glucose
Response to peptide hormone
Spindle organization
Regulation of cytokinesis
Regulation of G2/M transition of mitotic...
Mitotic sister chromatid segregation
Negative regulation of telomere maintena...
Cellular oxidant detoxification
Keratinocyte migration
Microtubule bundle formation
Mitotic spindle organization
Exit from mitosis
Cytolysis
Positive regulation of ubiquitin protein...
Mitotic centrosome separation
Protein K27-linked ubiquitination
Ribosome-associated ubiquitin-dependent...
Rich Ratio
Qvalue
Gene Number
D
GSEA plot:Cell Cycle (LOVO-EGFR vs Control
NOM p-Value<0.0001
E
84 cell cycle genes heatmap
high
low
Control OE-EGFR
F
WT
E545K
H1047R
Vehicle
Pal
EGFR
G
Vehicle
Pal
ns
***
*
WT
E545K
H1047R
H
Patient's data-derived palbociclib resistant genes (Dataset analyzed:GSE186901. Selection criterion: Expression alteration |log2FC|>0.5 in patients with disease progression after palbociclib treatment.)
+
Literature-based palbociclib resistance genes
Palbociclib resistant gene set (1716 genes)
I
GSEA plot: Palbociclib resistant genes set (PIK3CA MUT±S-CM)
NOM p-Value=0.001
GSEA plot: Palbociclib resistant genes set (OE-EGFR vs Control)
NOM p-Value=0.024
J
LOVO
SW620
Control
OE-EGFR
RB 110kDa
p-RB 110kDa
WEE1 95kDa
PKMYT1 60kDa
p16 16kDa
p21 21kDa
β-actin 44kDa
K
LOVO
SW620
Control
OE-EGFR
EGFR 175kDa
p-EGFR 175kDa
pAKT-308 60kDa
pAKT-473 60kDa
CDK2 33kDa
CDK4 36kDa
CDK6 30kDa
β-actin 44kDa

**Figure S3. SASP factors in palbociclib-induced PIK3CA mutant cell lines lead to resistance by upregulating EGFR.**

(**A**) KEGG analysis of RNA-sequencing datas from LOVO-PIK3CA MUT cells with S-CM treatment. (**B**) GSEA enrichment analysis of RNA-sequencing data from LOVO-PIK3CA MUT cells with S-CM treatment. (**C**) GO analysis of mass spectrometry differentially changed proteins after LOVO stable overexpression of EGFR. (**D**) GSEA enrichment analysis of Cell Cycle pathways based on mass spectrometry datas of LOVO stable overexpression of EGFR. (**H**) Heatmap indicating the expression of 84 cell cycle genes after overexpressed EGFR gene in LOVO cells. (**F**) Immunohistochemical analysis of EGFR expression in LOVO-PIK3CA WT and MUT cells mouse CDX models following palbociclib treatment. Quantification from image (**F**) is shown in **G**. (**H**) Schematic representation of the palbociclib resistance gene set. (**I**) GSEA enrichment analysis of S-CM and OE-EGFR differentially expressed genes and proteins in the palbociclib resistance gene set. (**J**) Immunoblot for palbociclib resistance proteins expression from LOVO and SW620 cell lines after overexpressed EGFR gene. (**K**) Immunoblot for cell cycle protein expression from LOVO and SW620 cell lines after overexpressed EGFR gene. Mean ± SD; *, $P < 0.05$, **, $P < 0.01$; and ***, $P < 0.001$.

Fig. S4

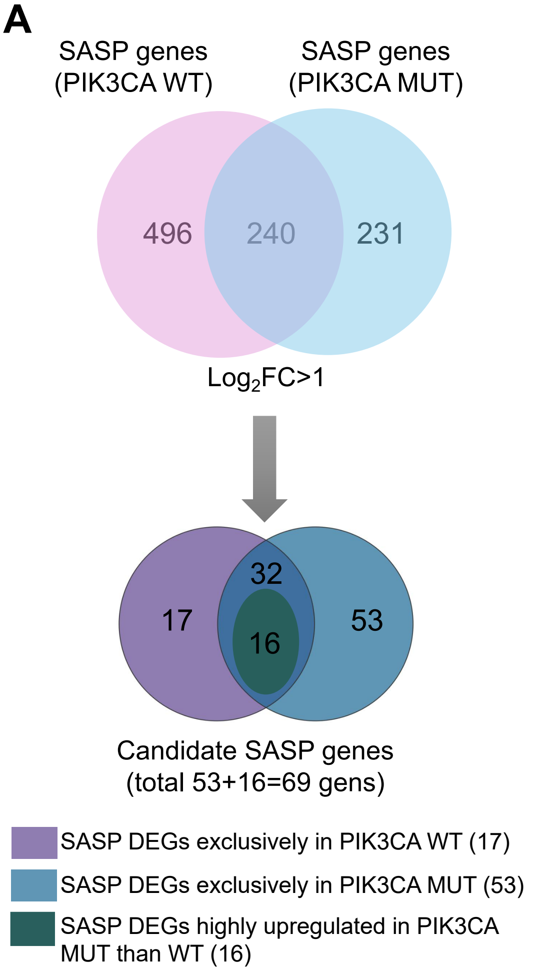


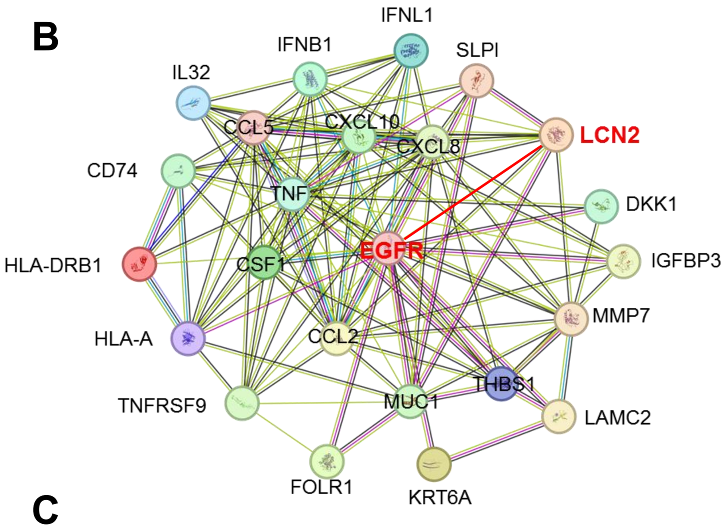


C

| | node1 | node2 | combined score |
|---|---|---|---|
| 1 | EGFR | MUC1 | 0.999 |
| 2 | EGFR | IGFBP3 | 0.981 |
| 3 | EGFR | CSF1 | 0.941 |
| 4 | EGFR | TNF | 0.89 |
| 5 | EGFR | IFNB1 | 0.877 |
| 6 | EGFR | CXCL8 | 0.792 |
| 7 | **EGFR** | **LCN2** | **0.729** |
| 8 | EGFR | CCL2 | 0.725 |
| 9 | EGFR | LAMC2 | 0.722 |
| 10 | EGFR | FOLR1 | 0.721 |
| 11 | EGFR | TNFRSF9 | 0.718 |
| 12 | EGFR | THBS1 | 0.708 |

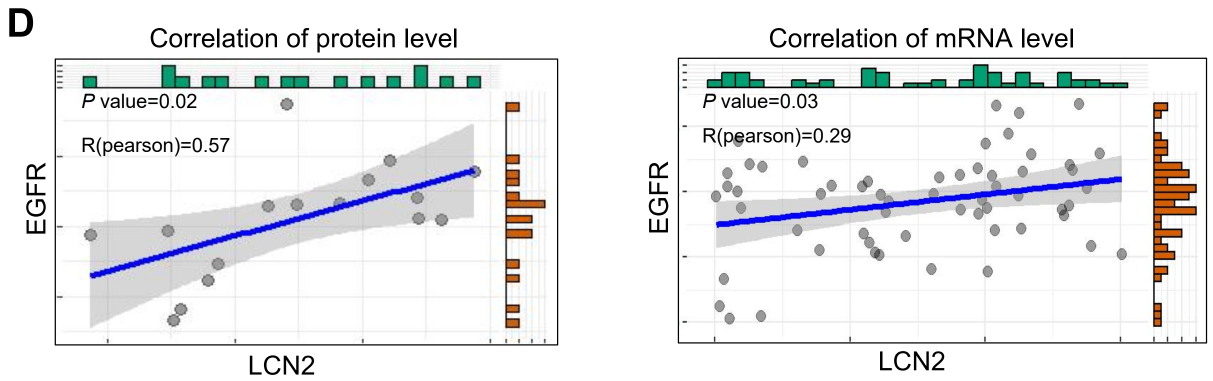


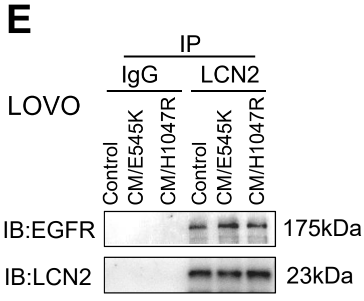


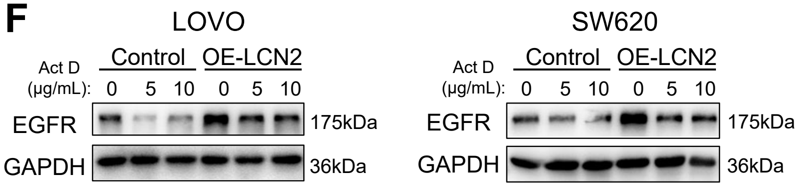

**Figure S4. The SASP factor LCN2 enhances the expression of EGFR by suppressing ubiquitination and lysosomal degradation pathways.**

(**A**) Schematic diagram of screening significantly expressed SASP genes in LOVO-PIK3CA WT and LOVO-PIK3CA MUT cell lines with palbociclib treatment. (**B**) The STRING website analysis results showed 23 SASP factors that interacted with EGFR protein. **C,** Protein interaction score list display from **B**. (**D**) Protein and mRNA level analysis of correlation data between LCN2 and EGFR comes from the CCLE database. (**E**) Co-IP immunoblot of EGFR-Flag and LCN2-Flag in transfected HEK293T cells under S-CM treatment. (**F**) Immunoblot analysis of EGFR protein expression in Act D time gradient treated LOVO and SW620 cells after overexpression of LCN2 gene.

Fig. S5

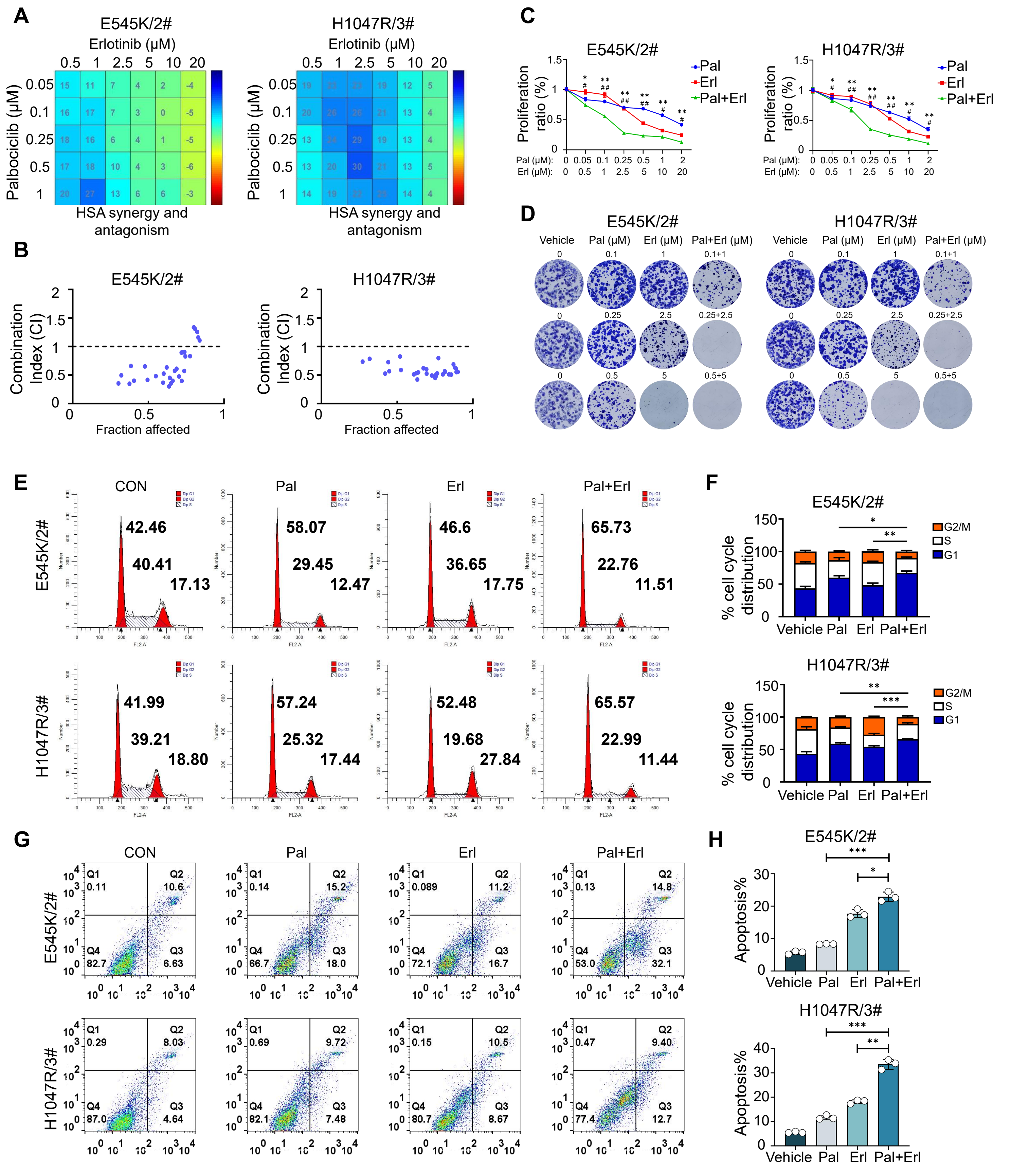
A
E545K/2#
H1047R/3#
Erlotinib (μM)
Palbociclib (μM)
HSA synergy and antagonism
B
Combination Index (CI)
Fraction affected
C
Proliferation ratio (%)
Pal
Erl
Pal+Erl
Pal (μM):
Erl (μM):
D
Vehicle
Pal (μM)
Erl (μM)
Pal+Erl (μM)
E
CON
Pal
Erl
Pal+Erl
42.46
40.41
17.13
58.07
29.45
12.47
46.6
36.65
17.75
65.73
22.76
11.51
41.99
39.21
18.80
57.24
25.32
17.44
52.48
19.68
27.84
65.57
22.99
11.44
F
% cell cycle distribution
G2/M
S
G1
Vehicle
G
Q1
Q2
Q3
Q4
H
Apoptosis%

**Figure S5. The screening of a drug library for the combination of erlotinib and palbociclib reveals a significant synergistic anti-tumor effect.**

(**A**) HSA synergy matrices showing the interaction between palbociclib and erlotinib in E545K/2# and H1047R/3# cell lines (HSA > 0 indicate synergistic effects). (**B**) CI plots showing the sturdy synergistic effect of palbociclib and erlotinib in E545K/2# and H1047R/3# cell lines (CI < 1 indicates a synergistic effects). (**C**) Growth inhibition curves of E545K/2# and H1047R/3# cell lines treated with fixed concentration-combinations of palbociclib and erlotinib. (**D**) Images showing the colony formation of E545K/2# and H1047R/3# cell lines treated with palbociclib and/or erlotinib. (**E**) Cell cycle analysis of E545K/2# and H1047R/3# cell lines upon treatment with palbociclib or erlotinib alone or in combination. (**F**) The quantification of cell fractions in G1, S and G2/M phases in **E**. (**G**) Apoptosis analysis of E545K/2# and H1047R/3# cell lines upon treatment of palbociclib and/or erlotinib. Quantification of apoptotic cells in (**H**) by image (**G**). Mean ± SD; *, $P < 0.05$, **, $P < 0.01$; and ***, $P < 0.001$.

Fig. S6

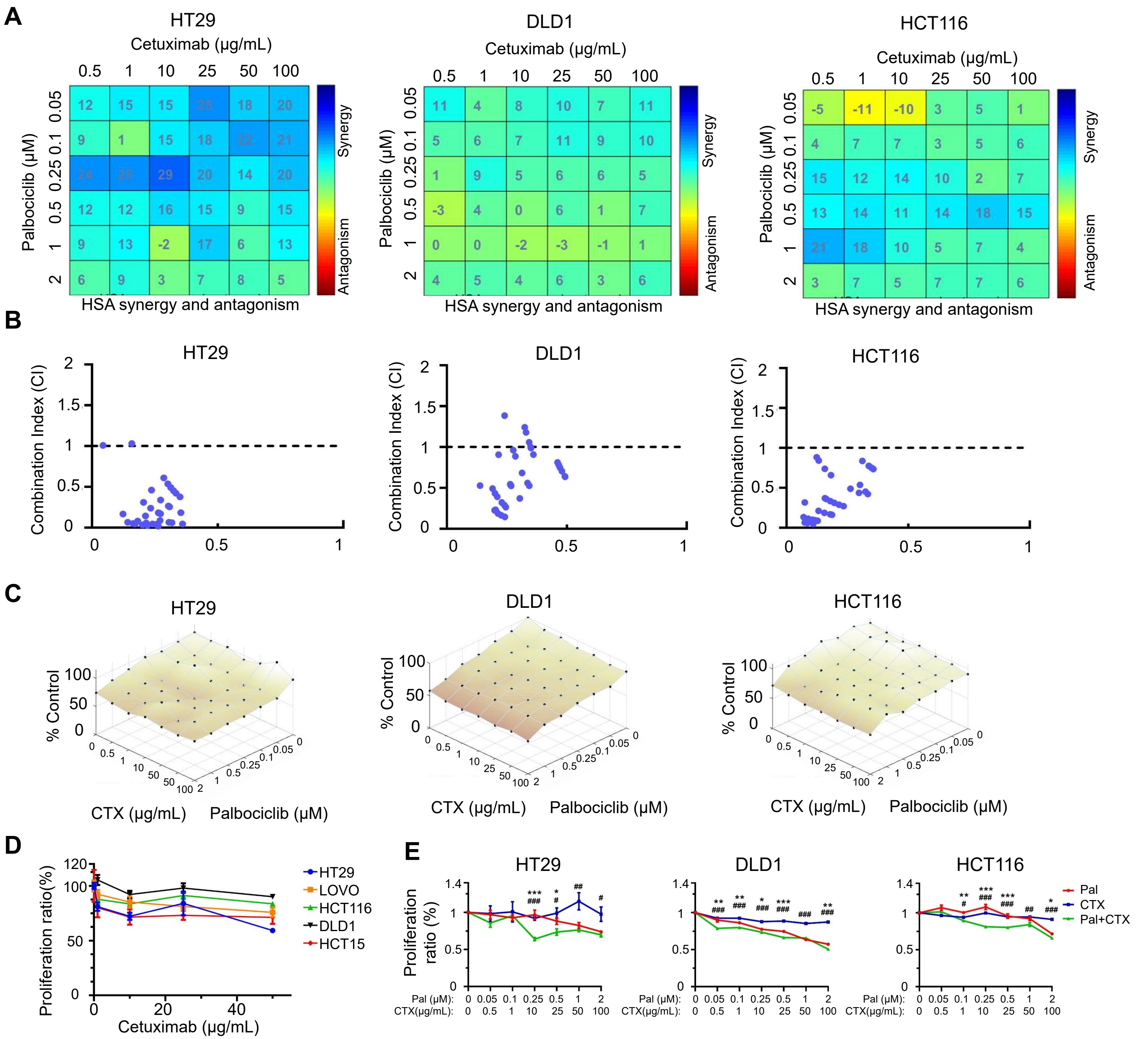
A
HT29
Cetuximab (μg/mL)
0.5 1 10 25 50 100
Palbociclib (μM)
0.05 0.1 0.25 0.5 1 2
12 15 15 25 18 20
9 1 15 18 22 21
24 25 29 20 14 20
12 12 16 15 9 15
9 13 -2 17 6 13
6 9 3 7 8 5
Synergy
Antagonism
HSA synergy and antagonism
DLD1
Cetuximab (μg/mL)
0.5 1 10 25 50 100
Palbociclib (μM)
0.05 0.1 0.25 0.5 1 2
11 4 8 10 7 11
5 6 7 11 9 10
1 9 5 6 6 5
-3 4 0 6 1 7
0 0 -2 -3 -1 1
4 5 4 6 3 6
Synergy
Antagonism
HSA synergy and antagonism
HCT116
Cetuximab (μg/mL)
0.5 1 10 25 50 100
Palbociclib (μM)
0.05 0.1 0.25 0.5 1 2
-5 -11 -10 3 5 1
4 7 7 3 5 6
15 12 14 10 2 7
13 14 11 14 18 15
21 18 10 5 7 4
3 7 5 7 7 6
Synergy
Antagonism
HSA synergy and antagonism
B
HT29
DLD1
HCT116
Combination Index (CI)
0 0.5 1 1.5 2
0 0.5 1
C
HT29
DLD1
HCT116
% Control
0 50 100
CTX (μg/mL)
Palbociclib (μM)
D
Proliferation ratio(%)
0 50 100 120
0 20 40
Cetuximab (μg/mL)
HT29
LOVO
HCT116
DLD1
HCT15
E
HT29
DLD1
HCT116
Proliferation ratio (%)
0 0.5 1 1.4
Pal (μM): 0 0.05 0.1 0.25 0.5 1 2
CTX(μg/mL): 0 0.5 1 10 25 50 100
Pal
CTX
Pal+CTX

**Figure S6. Cetuximab in combination with palbociclib exhibits a moderate anti-tumor efficacy.**

(**A**) HSA synergy matrices showing the interaction between palbociclib and cetuximab in HT29, DLD1 and HCT15 cell lines (HSA > 0 indicate synergistic effects). (**B**) CI plots showing the sturdy synergistic effect of palbociclib and erlotinib in HT29, DLD1 and HCT15 cell lines (CI < 1 indicates a synergistic effects). (**C**) Growth inhibition curves of HT29, DLD1 and HCT15 cell lines treated with cetuximab. (**D**) Growth inhibition curves of HT29, DLD1 and HCT15 cell lines treated with fixed concentration-combinations of palbociclib and cetuximab. (**E**) Surface plots generated by dose-matrices of the proliferation rates up treatment of palbociclib and cetuximab at different concentrations. (**F**) Images showing the colony formation of of HT29, DLD1 and HCT15 cell lines treated with palbociclib and/or cetuximab. Mean ± SD; *, $P < 0.05$, **, $P < 0.01$; and ***, $P < 0.001$.

Fig. S7

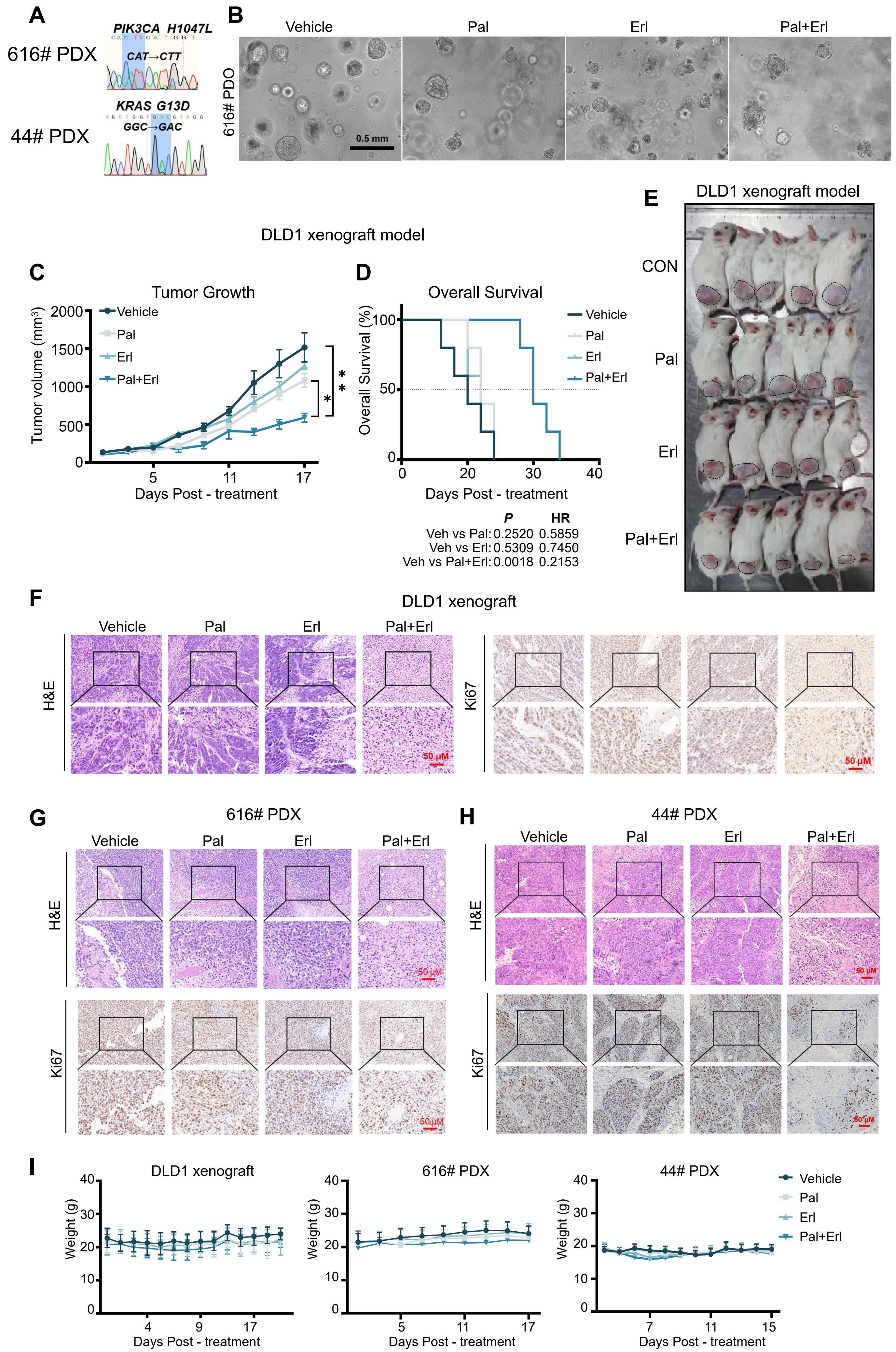
A
616# PDX
PIK3CA H1047L
CAT→CTT
44# PDX
KRAS G13D
GGC→GAC
B
Vehicle
Pal
Erl
Pal+Erl
616# PDO
0.5 mm
DLD1 xenograft model
C
Tumor Growth
Tumor volume (mm³)
Vehicle
Pal
Erl
Pal+Erl
Days Post - treatment
D
Overall Survival
Overall Survival (%)
Days Post - treatment
P HR
Veh vs Pal: 0.2520 0.5859
Veh vs Erl: 0.5309 0.7450
Veh vs Pal+Erl: 0.0018 0.2153
E
DLD1 xenograft model
CON
Pal
Erl
Pal+Erl
F
DLD1 xenograft
H&E
Ki67
50 μM
G
616# PDX
H
44# PDX
I
DLD1 xenograft
616# PDX
44# PDX
Weight (g)
Days Post - treatment

**Figure S7. PE-combination exhibited significant efficacy in PDO and CDX models with PIK3CA mutation.**

(**A**) Sequencing identified the H1047L mutation in the PIK3CA gene and the G13D mutation in the KRAS gene in PDX models 616# and 44#. (**B**) Representative images of PDOs (derived from 616#) treated with palbociclib (1 μM) alone or in combination with erlotinib (10 μM). Scale bar, 0.5 mm. (**C**) Tumor volume curves of CDX (DLD1) after palbociclib and/or erlotinib treatment. (**D**) Kaplan-Meier curves depicting OS for CDX (DLD1) mice treated with palbociclib and/or erlotinib. (**E**) Representative images of CDX (DLD1) mice upon treatment of palbociclib and/or erlotinib. (**F-H**) Tumor tissue stained with H&E and Ki-67 of CDX (DLD1) and PDX (616#, 44#) upon treatment of palbociclib and/or erlotinib (scale bar: 50 μm). (**I**) The curves of CDX (DLD1) and PDX (616#, 44#) body weights upon treatment of palbociclib and/or erlotinib. Mean ± SD; Mean ± SEM; *, $P < 0.05$, **, $P < 0.01$; and ***, $P < 0.001$.
