## Supplementary material for "Oncogene-induced senescence and senescence-associated secretory phenotype (SASP) determine the efficacy of palbociclib in PIK3CA mutated colorectal cancer": Palbociclib resistance genes from literature

**Table1. Palbociclib resistance genes from literature：**

| pathway&gene | gene name | condition |
|---|---|---|
| Cell cycle-specific mechanisms | MDM2[1] | overexpression |
| | MDM4[2] | overexpression |
| | WEE1[3] | overexpression |
| | CCNE1/2[4] | amplification |
| | CDK2[5] | amplification |
| | CDK6[6] | amplification |
| | CDKN2A[7, 8] | mutations |
| | CDKN2B[9] | mutations |
| | RB1[10] | loss |
| | E2F[4] | amplification |
| | CDK4[11] | amplification |
| | CDK7[12] | overexpression |
| | HDAC[13] | loss |
| | FZR1[14] | loss |
| | FGFR1[15] | amplification |
| Activation of the FGFR pathway | FGF2[16] | amplification |
| | ER[17] | loss |
| Activation of the PI3K/AKT/mTOR pathway | AKT1[18] | overexpression |
| | AKT3[18] | overexpression |
| | PIK3CA[18] | mutation |
| | ESR1[19] | mutation |
| | PDK1[20] | overexpression |
| | FOXM1[21] | overexpression |
| TP53 | TP53[22] | mutation |
| HER2 | HER2[23] | overexpression |
| AURKA | AURKA[24] | mutation/amplification |
| PLK1 | PLK1[25] | overexpression |
| S6K1 | S6K1[26] | amplification |

### REFERENCES：


1. Laroche-Clary A *et al.* Combined targeting of MDM2 and CDK4 is synergistic in dedifferentiated liposarcomas. *J Hematol Oncol*. **10**, (2017).
2. AbuHammad S *et al.* Regulation of PRMT5-MDM4 axis is critical in the response to CDK4/6 inhibitors in melanoma (vol 116, pg 17990, 2019). *P Natl Acad Sci USA*. **117**, 9644-9645 (2020).
3. Matheson CJ, Backos DS, Reigan P. Targeting WEE1 Kinase in Cancer. *Trends Pharmacol Sci*. **37**, 872-881 (2016).
4. Taylor-Harding B *et al.* Cyclin E1 and RTK/RAS signaling drive CDK inhibitor resistance via activation of E2F and ETS. *Oncotarget*. **6**, 696-714 (2015).
5. Patel P *et al.* Dual Inhibition of CDK4 and CDK2 via Targeting p27 Tyrosine Phosphorylation Induces a Potent and Durable Response in Breast Cancer Cells. *Molecular Cancer Research*. **16**, 361-377 (2018).

6. Yang C *et al.* Acquired CDK6 amplification promotes breast cancer resistance to CDK4/6 inhibitors and loss of ER signaling and dependence. *Oncogene*. **36**, 2255-2264 (2017).
7. Cen L *et al.* p16-Cdk4-Rb axis controls sensitivity to a cyclin-dependent kinase inhibitor PD0332991 in glioblastoma xenograft cells. *Neuro-Oncology*. **14**, 870-881 (2012).
8. Liu Y *et al.* p16(INK4a) expression in retinoblastoma: a marker of differentiation grade. *Diagn Pathol*. **9**, 180 (2014).
9. Green JL *et al.* Direct CDKN2 Modulation of CDK4 Alters Target Engagement of CDK4 Inhibitor Drugs. *Mol Cancer Ther*. **18**, 771-779 (2019).
10. Condorelli R *et al.* Polyclonal RB1 mutations and acquired resistance to CDK 4/6 inhibitors in patients with metastatic breast cancer. *Ann Oncol*. **29**, 640-645 (2018).
11. Olanich ME *et al.* CDK4 amplification reduces sensitivity to CDK4/6 inhibition in fusion-positive rhabdomyosarcoma. *Cancer Res*. **75**, (2015).
12. Schachter MM *et al.* A Cdk7-Cdk4 T-loop phosphorylation cascade promotes G1 progression. *Mol Cell*. **50**, 250-260 (2013).
13. Zhou Y *et al.* HDAC5 Loss Impairs RB Repression of Pro-Oncogenic Genes and Confers CDK4/6 Inhibitor Resistance in Cancer. *Cancer Res*. **81**, 1486-1499 (2021).
14. The I *et al.* Rb and FZR1/Cdh1 determine CDK4/6-cyclin D requirement in C. elegans and human cancer cells. *Nature Communications*. **6**, (2015).
15. Mao PP, Kusiel J, Cohen O, Wagle N. The role of FGF/FGFR axis in resistance to SERDs and CDK4/6 inhibitors in ER plus breast cancer. *Cancer Res*. **78**, (2018).
16. Formisano L *et al.* Aberrant FGFR signaling mediates resistance to CDK4/6 inhibitors in ER+ breast cancer. *Nat Commun*. **10**, 1373 (2019).
17. Finn RS, Aleshin A, Slamon DJ. Targeting the cyclin-dependent kinases (CDK) 4/6 in estrogen receptor-positive breast cancers. *Breast Cancer Res*. **18**, 17 (2016).
18. Takeshita T *et al.* Clinical significance of plasma cell-free DNA mutations in PIK3CA, AKT1, and ESR1 gene according to treatment lines in ER-positive breast cancer. *Mol Cancer*. **17**, 67 (2018).
19. Gyanchandani R *et al.* Detection of ESR1 mutations in circulating cell-free DNA from patients with metastatic breast cancer treated with palbociclib and letrozole. *Oncotarget*. **8**, 66901-66911 (2017).
20. Jansen VM *et al.* Kinome-Wide RNA Interference Screen Reveals a Role for PDK1 in Acquired Resistance to CDK4/6 Inhibition in ER-Positive Breast Cancer. *Cancer Res*. **77**, 2488-2499 (2017).
21. Anders L *et al.* A systematic screen for CDK4/6 substrates links FOXM1 phosphorylation to senescence suppression in cancer cells. *Cancer Cell*. **20**, 620-634 (2011).
22. Bellutti F *et al.* CDK6 Antagonizes p53-Induced Responses during Tumorigenesis. *Cancer Discov*. **8**, 884-897 (2018).
23. Goel S *et al.* Overcoming Therapeutic Resistance in HER2-Positive Breast Cancers with CDK4/6 Inhibitors. *Cancer Cell*. **29**, 255-269 (2016).
24. Lee S *et al.* Exploratory analysis of biomarkers associated with clinical outcomes from the study of palbociclib plus endocrine therapy in premenopausal women with hormone receptor-positive, HER2-negative metastatic breast cancer. *Breast*. **62**, 52-60 (2022).
25. Liang H *et al.* Targeting CBX3 with a Dual BET/PLK1 Inhibitor Enhances the Antitumor Efficacy of CDK4/6 Inhibitors in Prostate Cancer. *Adv Sci (Weinh)*. **10**, e2302368 (2023).
26. Mo H *et al.* S6K1 amplification confers innate resistance to CDK4/6 inhibitors through activating c-Myc pathway in patients with estrogen receptor-positive breast cancer. *Mol Cancer*. **21**, 171 (2022).
